# Quantifying sporozoite inoculum dynamics with the SpitGrid reveals temporal decay and behavioral determinants of *Plasmodium* transmission

**DOI:** 10.64898/2026.01.06.697895

**Authors:** Felix Evers, Geert-Jan van Gemert, Kjerstin Lanke, Wouter Graumans, Chiara Andolina, Benjamin Mordmüller, Teun Bousema, Felix JH Hol

## Abstract

The number of *Plasmodium* sporozoites expelled by an infected mosquito during a bite, the inoculum size, is a critical but rarely measured factor in transmission. Its temporal dynamics, individual variation, and behavioral underpinnings remain poorly understood. To address this, we developed the SpitGrid, a modular platform that supports natural probing and feeding on artificial substrates, while enabling behavioral characterization combined with non-lethal, sensitive measurement of inoculum size for individual mosquitoes in high-throughput. Using the SpitGrid, we determined inoculum size of *Plasmodium falciparum*-infected *Anopheles stephensi* mosquitoes over time and identified a robust temporal structure. Inoculum size peaked shortly after the extrinsic incubation period, and declined approximately log-linearly post peak. Salivary gland parasite load decreased only moderately and was positively associated with inoculum size during the early phase of sporozoite expelling, yet explained only part of the variation observed in inoculum size. Fine-grained behavioral classification confirmed that probing alone can deliver substantial inocula, and revealed that probing more often increased inoculum size while the duration of probing, or if this led to feeding, had little effect. Application of the SpitGrid to *Plasmodium vivax*-infected mosquitoes similarly showed a temporally structured expelling profile, with generally smaller inocula and a higher proportion of non-expelling mosquitoes. Our observations reveal that *Plasmodium* infectivity in mosquitoes is a highly dynamics process, challenging the assumption that per-bite transmission potential remains constant over time. We establish the SpitGrid as a flexible tool to gain a nuanced understanding of the determinants of transmission, refine epidemiological models, and evaluate the efficacy of vector- based interventions under biologically relevant conditions.

**Highlights:** We develop the SpitGrid, a flexible, accessible, low-cost and animal free platform to measure sporozoite expelling

We identify a peak in *P. falciparum* and *P. vivax* sporozoite expelling by infected *Anopheles stephensi* mosquitoes early after completion of the extrinsic incubation period, followed by a gradual decline

We confirm a correlation between salivary gland load and inoculum size that weakens over time, indicating that a smaller fraction of the remaining salivary gland load is being expelled by older mosquitoes

We confirm that mosquito probing alone can deliver substantial sporozoite inoculum independently of a successful bloodmeal

By dissecting the behavioral parameters that determine inoculum size, we observe that the initiation of probing is a major determinant of inoculum size with relatively minor roles for probing time and whether biting results in engorgement

Compared to *P. falciparum*, we observe comparatively smaller inocula, in *P. vivax*-infected mosquitoes a higher rate of non-expellers and similar albeit more moderate temporal structure with an earlier onset of expelling, illustrating cross-species utility of the SpitGrid

## Introduction

Malaria transmission happens one bite at a time. With each bite, an infected mosquito can deliver *Plasmodium* sporozoites into the skin, initiating infection. The size of this inoculum is shaped by a variety of factors. From parasite quantity, development, and availability within the salivary glands to the dynamics of the mosquito bite and the physiology of an infected salivary apparatus, there are many plausibly important parameters that might determine inoculum size. As a result of this complexity, inoculum size is challenging to approximate in theory, and in practice outcomes of an infected mosquito bite are notoriously heterogenous^1–3^. Nevertheless, a nuanced understanding of the determinants of inoculum size is needed to move from simplistic binary infectiousness assumptions toward a biologically grounded understanding that allows quantitative assessment of transmission, and its disruption.

More than a century ago, Ronald Ross, who originally discovered that malaria parasites are transmitted by mosquitoes, already pondered the determinants of the inoculum. He speculated that it must depend on the “number of spores in the insect’s glands” and the “number of times the insect bites it victim” as mosquitoes “inject their poison before commencing to suck blood”^4^. In 1945 Percy Shute investigated whether repeated feeds deplete parasites from the salivary glands, and suggested that bites delivered shortly after salivary gland invasion may inject far more sporozoites than later bites^5^. But those ideas were difficult to test and have received limited attention since then. Half a century later, researchers investigated the inoculum directly using forced salivation of individual mosquitoes estimating median inoculum size around ten sporozoites^6–10^, far lower than historical estimates^5^ and 2-3 orders of magnitude removed from what is administered to reliably induce malaria in human challenge models^11^. Recently, more direct methods have been pioneered in which infected mosquitoes feed on the ear of anesthetized mice^12^ or a blood-soaked collagen matrix^13^, and estimates of a median inoculum moved up to around 1000 sporozoites per bite. These advances enabled a more nuanced understanding of the sporozoite inoculum, and how it relates to infection status of the mosquito and provided solid evidence that salivary gland sporozoite load is positively associated with inoculum size. At the same time, the labor intensiveness, high cost, and/or use of animals have thus far limited widespread use of these methods. As a result, questions surrounding the effect of interventions, age of infection or mosquitoes, different parasite vector combinations, or variations in biting behavior on inoculum size all remain unexplored. This gap is particularly pronounced for *Plasmodium vivax*, where experimental infections are scarce and basic features such as inoculum size and the temporal onset of transmissibility remain essentially unaddressed.

In practice, *Plasmodium* transmission is generally assessed by oocyst prevalence or density in the mosquito midgut or, less commonly, the quantity of sporozoites in the salivary glands. While the latter has been shown to be positively correlated with the size of the inoculum^12, 13^ (and infectivity^14^), assessing it is invariably fatal to the mosquito and does not explain most of the variation in inoculum size. It furthermore does not capture mosquito biology nor behavior. This is important because mosquito biting is not a binary act: length of probing or feeding, variability in salivation volume, successive probing bouts, and the decision to engorge or disengage all vary within and between individual mosquitoes and may profoundly impact sporozoite transmission. Interestingly, insecticide resistance and sublethal exposure to endectocides both significantly alter blood feeding behavior^15–18^ yet it remains unknown if this affects sporozoite transmission capacity.

Despite abundant evidence that biting behavior is plastic, the direct impact of this plasticity on transmission has been under-explored and is often collapsed into a one-size-fits-all “bite” event in epidemiological transmission models. Transmission models furthermore generally assume that every mosquito, once the extrinsic incubation period (EIP) has passed, can transmit *Plasmodium* with every subsequent bloodmeal until death, with a fixed probability per bite. This does not only neglect heterogeneity between mosquito bites but also assumes that transmission of parasites, all else being equal, is static over the lifetime of a mosquito. The data underlying this assumption, however, is sparse and mixed. Some (historic) work has suggested a decline in infectivity over time while other work has suggested no drop in expelled sporozoites and infection status in the first week after EIP has elapsed^5, 12, 19^. To address this we set out to measure the inoculum size, directly from the bite, over time while preserving and recording the mechanics of probing and engorgement. To enable this, we developed the SpitGrid, a platform that combines an easily reproducible and scalable grid of individually housed mosquitoes, an artificial bite substrate allowing video recording and behavioral classification of mosquitoes, and a non- destructive qPCR-based readout for accurate determination of inoculum size.

Applying this approach, we find robust evidence for both *P. falciparum* and *P. vivax* that there is a relatively short peak transmission window at which high numbers of sporozoites are expelled, that is followed by a rapid decay in inoculum size over time. We also find indications that inoculum size is lower for *P. vivax* compared to *P. falciparum*. We further delineate how probing and feeding behaviors impact inoculum size. Taken together, its accessibility, flexibility and high throughput make the SpitGrid a powerful platform to quantify sporozoite transmission dynamics of different *Plasmodium* and *Anopheles* species over time and evaluate interventions that shift when and how many parasites are released.

## Results

### The SpitGrid enables direct and non-fatal assessment of sporozoite expelling over time

The SpitGrid was inspired by the observation that mosquitoes will readily probe and feed from an artificial substrate if specific requirements are met^20^. Skin-like temperature, elevated CO_2_, ATP, NaCl, sodium bicarbonate, neutral pH, low concentration glucose and a pierceable membrane all positively influence the likelihood of anopheline mosquitoes to engage in blood feeding behavior on an artifcial substrate^21, 22^. We have found that this holds true when the artificial substrate is presented in form of an agarose hydrogel, which obviates the need for a membrane cover and facilitates downstream DNA or RNA isolation. To assay sporozoite expelling in a scalable high-throughput format, a gridded cage consisting of 4x6 individual cells was constructed from transparent acrylic (Fig. 1A). A perforated bottom in each cell facilitates mosquito access to an artificial bite substrate. A tiled array of artificial bite substrates was arranged on a grid corresponding to the cage grid and warmed to skin temperature using a controllable transparent heater. A camera monitored behavior and feeding status of mosquitoes in each individual grid cell. Mosquitoes were allowed to probe and feed on an artifical bite substrate (Fig. 1B) until cessation of activity up to a maximum of thirty minutes. Subsequently, artificial bite substrates were collected for quantification of deposited sporozoites through qPCR (see Methods). On any given experiment day, approximately 50% of mosquitoes interacted with the bite substrate, either only probing, or also engorging on artificial blood meal. As feeding on the artificial blood meal does not induce egg development, it does not incur the multi-day refractory period typically following blood feeding^23^. Short term cessation of blood feeding through abdominal stretch receptors still terminates blood feeding and subsequent seeking behavior as with a natural bloodmeal^24, 25^ but mosquitoes appear to appear to return baseline physiology within around 24 hours. This property can be exploited to follow repeated expelling (e.g. every two days) of individual mosquitoes after their infectious bloodmeal (dpi). This biting rhythm is highly relevant for the natural context in which anophelines often take multiple blood meals during a single gonotrophic cycle^26^.

**Figure 1.**
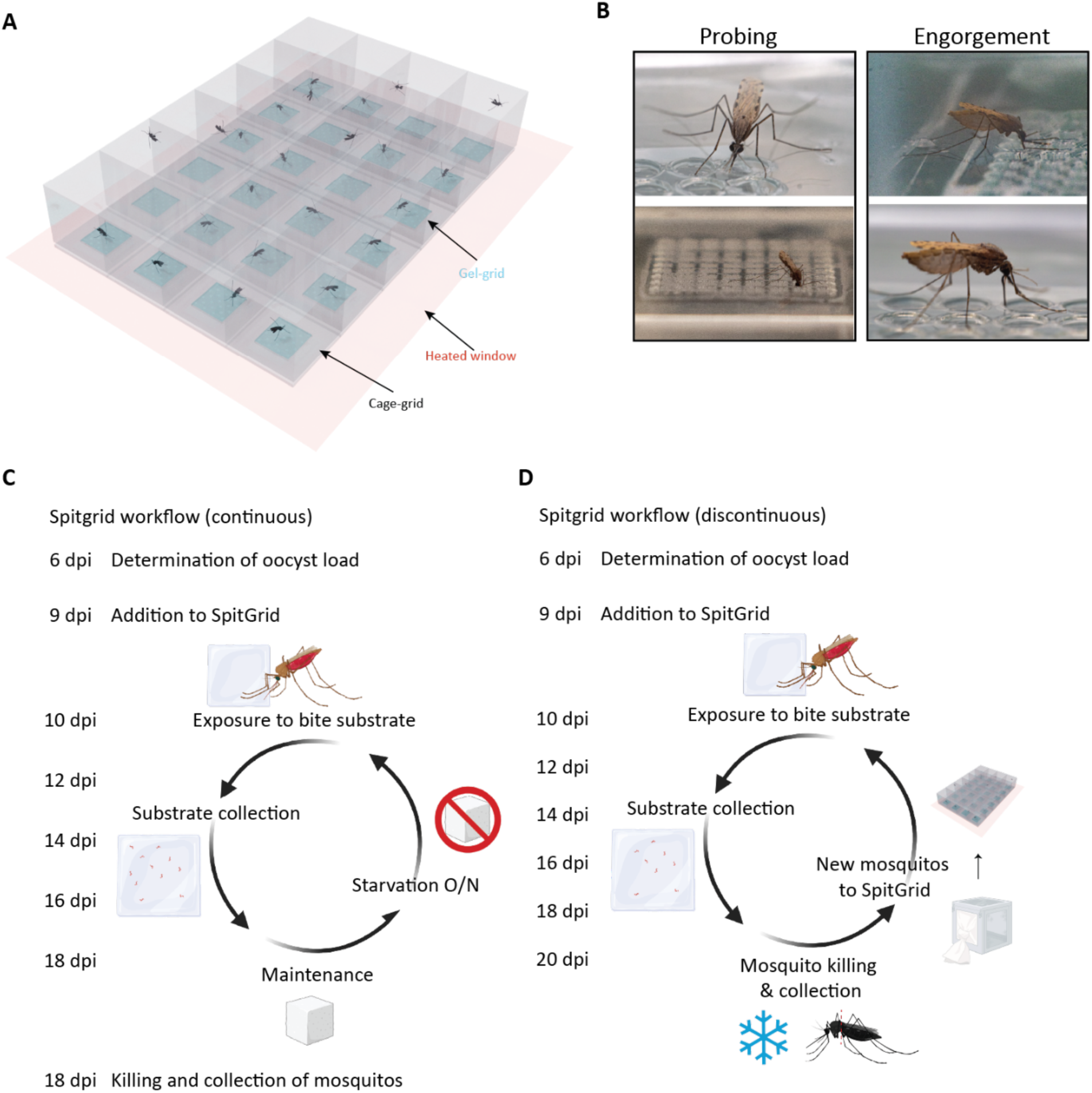
The SpitGrid **A** Schematic of the SpitGrid: a 4×6 acrylic cage-grid aligned to a gel-grid and mounted on a transparent glass heater to maintain the artificial bite substrate at skin-like temperature for an. A camera mounted below the heater records behavior. **B** Exemplary pictures of mosquitoes engaging in probing or engorgement in the SpitGrid. **C** Flowchart of the “continuous” implementation of the SpitGrid: Infected mosquitoes are introduced at 9 dpi (days post infection) and presented with the bite substrate on 10, 12, 14, 16 and 18 dpi while being maintained between sessions (continuous sugar access; starved overnight before each exposure). Parasite quantities in bite substrates and mosquitoes that died throughout or were killed at 18 dpi are determined through qPCR **D** Flowchart of the “discontinuous” implementation of the SpitGrid: new infected mosquitoes are added for each experimental day; after exposure mosquitoes are killed for thorax (salivary gland) qPCR.

### Sporozoite inoculum size is heterogenous and dynamic over time

We first deployed the SpitGrid in a discontinuous timecourse, in which mosquitoes were sacrificed and assessed for salivary gland load after each artificial bite substrate exposure (Fig. 1D). New mosquitoes were added to the SpitGrid for each time point, providing insight into the relationship between the inoculum deposited during a bite, and the mosquitoes’ salivary gland load at that time. In line with the expected extrinsic incubation period (EIP) of *P. falciparum*, we observed very little parasite expelling on 10 dpi. Over 80% of interacting mosquitoes expelled no sporozoites at all at this timepoint and the remainder expelled <100 sporozoites (Fig 2A, C). Corresponding salivary gland loads were low, but only 17% of mosquitoes was salivary gland negative at 10 dpi (Fig 2C). At 12 dpi salivary gland load and inocolum size had increased dramatically to their peak value (Fig. 2A). The fraction of mosquitos that probed or fed but did not expell sporozoites dropped from >80% at 10 dpi to ∼20% at 12 dpi and 43% of mosquitoes expelled more than 1,500 sporozoites (Fig. 2C). Furthermore, all salivary glands were positive with most in the range of 25,000 – 100,000 sporozoites (Fig. 2B). From 14 dpi onwards both salivary gland load and inoculum size decreased, but the decline differed in magnitude and shape (Fig. 2A). Salivary gland loads dropped to ∼25% of the peak value at 16 dpi, and plateaued thereafter. Inoculum size on the other hand declined continously until the experiment was terminated at 20 dpi at which point the median inoculum was only ∼ 3% the size of the peak value (Supp. Fig. 2C). As a result of the divergence of inoculum size and salivary gland load, the fraction of the salivary gland load that is expelled continously declined after peaking at 12 dpi (Fig. 2D). Remarkably the fraction of mosquitoes that failed to expell sporozoites remained relatively consistent at ∼20% despite all mosquitoes being salivary gland positive post 10 dpi (Fig. 2B,C). The decline in inoculum size appears to be driven by a decrease in mosquitoes that expell relatively high numbers of sporozoites (>500, Fig. 2C). A generalized linear model (GLM) with loglink and a single inflection at 12 dpi captured this pattern and confirms time as a significant predictor of incolum size both before (30 ± 11x / day) and after (0.63x ± 0.05 / day) 12 dpi (Supp. Fig 1A). A parallel model for the salivary gland load (Supp. Fig. 1B) showed a synchronized rise (2.5x ± 0.7 / day) to 12 dpi followed by a decline (0.83x ± 0.03 / day) thereafter (and a further slowdown in decline from 16 – 20 dpi). On all days, inoculum size spanned three orders of magnitude, which is a remarkable level of heterogeneity, especially since salivary gland loads showed considerably less variation (Fig. 2A). In summary, salivary gland load and particularly inoculum size are dynamic over time, showing an initial peak followed by subsequent decline.

**Figure 2.**
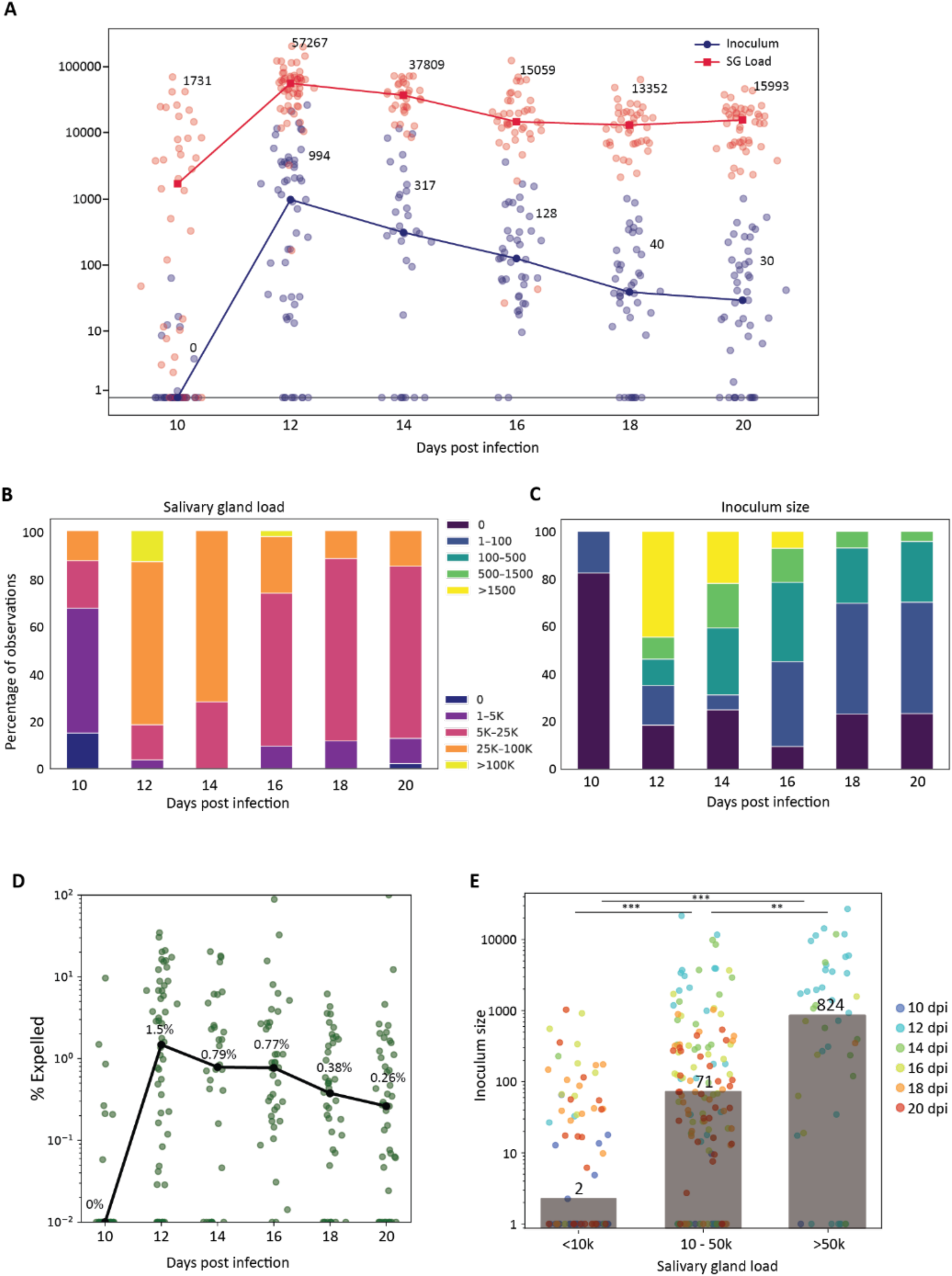
Inoculum size and salivary gland load are dynamic over time. **A** Median salivary gland load (SG load, red) and inoculum size (Inoculum, blue) from 10 dpi to 20 dpi. Both metrics show an initial increase, peak at 12 dpi, and subsequent decline. **B** Relative frequency of different salivary gland load ranges (0, 1-5000, 5001-25000, 25001-100000, >100000) over time (10 - 20 dpi). From 12 - 20 dpi a major shift from the 25000-100000 to the 5000-25000 bin can be observed. Underlying datapoints are shown as transparent dots **C** Relative frequency of different inoculum size ranges (0, 1-100, 101-500, 501-1500, >1500) over time (10 - 20 dpi). **D** Median percentage of parasites that are expelled relative to salivary gland load (black). Inoculum size relative to salivary gland load continuously decreases after an initial peak at 12 dpi. Underlying datapoints are shown as transparent dots **E** Median inoculum size of mosquitoes with different ranges of salivary gland load (<10000, 10000-50000, >50000). Ranges are based on thresholds suggested in Kanatani et al.^12^. Underlying datapoints are connected to timepoints (blue = 10 dpi, teal = 12 dpi, green = 14 dpi, khaki = 16 dpi, orange = 18 dpi, red = 20 dpi) and shown as transparent dots.

### The relationship between inoculum size and salivary gland load

We next investigated if salivary gland load predicts inocolum size. Across days, we observed a highly significant albeit modest positive relationship between salivary gland load and inoculum size, with salivary gland load predicting 21% of the variability in inoculum size (Supp Fig. 2A). This association appeared strong on the peak expelling day but weak or non-significant on other days (Supp. Fig. 2A). Prior research has suggested a threshold model of sporozoite expelling in which infectivty and sporozoite output increases substantially once a threshold of salivary gland load is passed^12–14^. Subdividing mosquitoes into these bins, we indeed observed that inoculum size differs signficantly between the three salivary gland load groups (Fig. 2E), with salivary gland loads of less than 10k sporozoites resulting in small inocula, salivary gland loads of 10-50k having intermediate sized inocula, and salivary gland loads over 50k having large inocula. As both inoculum size and salivary gland load are a covariate of time since infection and the relation between salivary gland load and inoculum appears most prominent on 12 dpi (Supp. Fig. 2A), we tested whether salivary gland load remains a meaningful predictor when accounting for dpi. Using the same log-link GLM but incorporating salivary gland load as a predictor variable we found that salivary gland load remained signficantly associated with inoculum size, with every 10x increase in salivary gland load doubling inoculum size (Supp. Fig. 2D).

Taken together, our data support the intuitive and previously shown^12, 13^ positive association between salivary gland load and inoculum size but also highlight that a large part of the variability in inoculum size remains unexplained. This implies that on the individual level expelling is a highly stochastic process influenced by a complex set of variables which we explore below.

### Expelling trajectories of individual mosquitoes do not show signs of depletion

Next, we investigated the dynamics of the infectious capacity of an individual mosquito over longer periods through a continous timecourse in which individual mosquitoes were maintained in the SpitGrid from 9 to 18 dpi while being exposed to a fresh bite substrate on even days (Fig. 1C). This allowed us to examine how multiple feeding or probing bouts throughout a mosquito’s lifetime affect sporozoite expelling. In the SpitGrid (or when presented with a life host), not all mosquitoes feed or interact at each opportunity. Across experiments and days, roughly half remain passive when presented with the artificial bite substrate (Table 2.). As a result, individual mosquitoes fed or probed on varying numbers of days, ranging from one to all five possible days (Fig. 3B). Following these expelling trajectories, we found many examples of mosquitoes that expelled at multiple or all opportunities (Fig. 3A). This suggests that sporozoites are not meaninfully depleted or replenished within two days. To further explore the depletion hypothesis we also checked whether past feeding or probing was negatively associated with future inoculum size. We found that there is no signficant difference in inoculum size between mosquitoes that had a history of activity (probing or feeding) compared to those that remained passive (Fig. 3C). Inoculum size and timing was largely consistent with the discontinous timecourse except that peak expelling shifted from 12 dpi to 14 dpi (Supp. Fig. 3A) demonstrating that the SpitGrid is viable for long-term and repeated assessment of individual mosquito expelling without obvious modulation of behavioral or expelling parameters. Taken together, our data supports the notion that expellable sporozoites are not meaningfully depleted with expelling opportunities in 48 hour intervals, also in the context of multiple feeding with up to five feeds accross 8 days.

**Figure 3.**
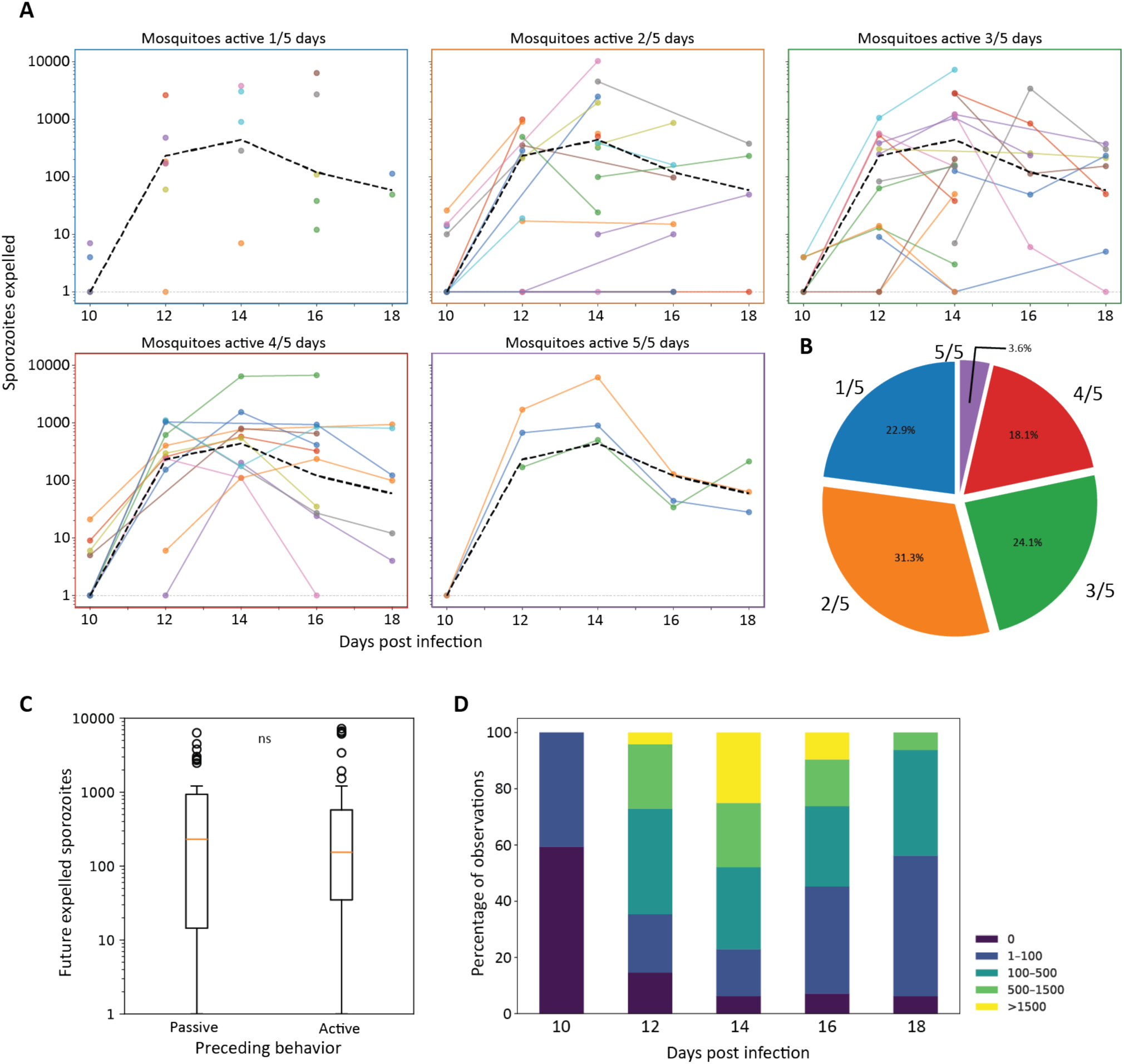
Past expelling does not decrease future expelling. **A** Inoculum size of mosquitoes that have fed on, or probed the artificial bite substrate 1, 2, 3, 4 or 5 out of 5 times. No systemic depletion after prior activity can be observed. Each color represents an individual mosquito with the constituting expelling events represented by colored circles. Median inoculum size of all mosquitoes over time is indicated with a black line. **B** Relative distribution of mosquitoes that have interacted 1 – 5 times **C** Boxplot of inoculum size for expelling events that were preceded by a passive day (no interaction with artificial bite substrate) or an active day (probing or feeding from artificial bite substrate). No significant differences were found (two-sided Mann-Whitney-U, *p* = 0.59) **D** Relative frequency of different inoculum size ranges (0, 1-100, 101-500, 501-1500, >1500) over time (10 – 18 dpi) in the continuous experiment.

**Table 1.** Discontinous timecourse experiment information Underlying mosquito cohort had an infection prevalence of 100% and an average 14.3 oocyst per midgut. Double/Single BM gives the ratio of interacting mosquitos that have received an additional uninfectious bloodmeal or not.

| Days post infection | Passive Mosquitoes | Feeding Mosquitoes | Probing Mosquitoes | Double / Single BM | Sample Size | Median SPZ Inoculum | SG Load |
| --- | --- | --- | --- | --- | --- | --- | --- |
| 10 | 56 (59%) | 12 (13%) | 26 (28%) | 0.90 | 38 | 0 | 1759 |
| 12 | 42 (44%) | 24 (25%) | 30 (31%) | 1.00 | 54 | 960 | 58141 |
| 14 | 64 (66%) | 16 (17%) | 16 (17%) | 0.68 | 32 | 317 | 37816 |
| 16 | 54 (57%) | 11 (12%) | 29 (31%) | 0.90 | 40 | 128 | 15060 |
| 18 | 53 (55%) | 8 (8%) | 34 (36%) | 0.83 | 42 | 40 | 13353 |
| 20 | 49 (51%) | 16 (17%) | 31 (32%) | 1.24 | 47 | 30 | 14969 |
| Average | 53 (55%) | 15 (16%) | 28 (29%) | 1 | 42 | 246 | 23516 |

**Table 2.**
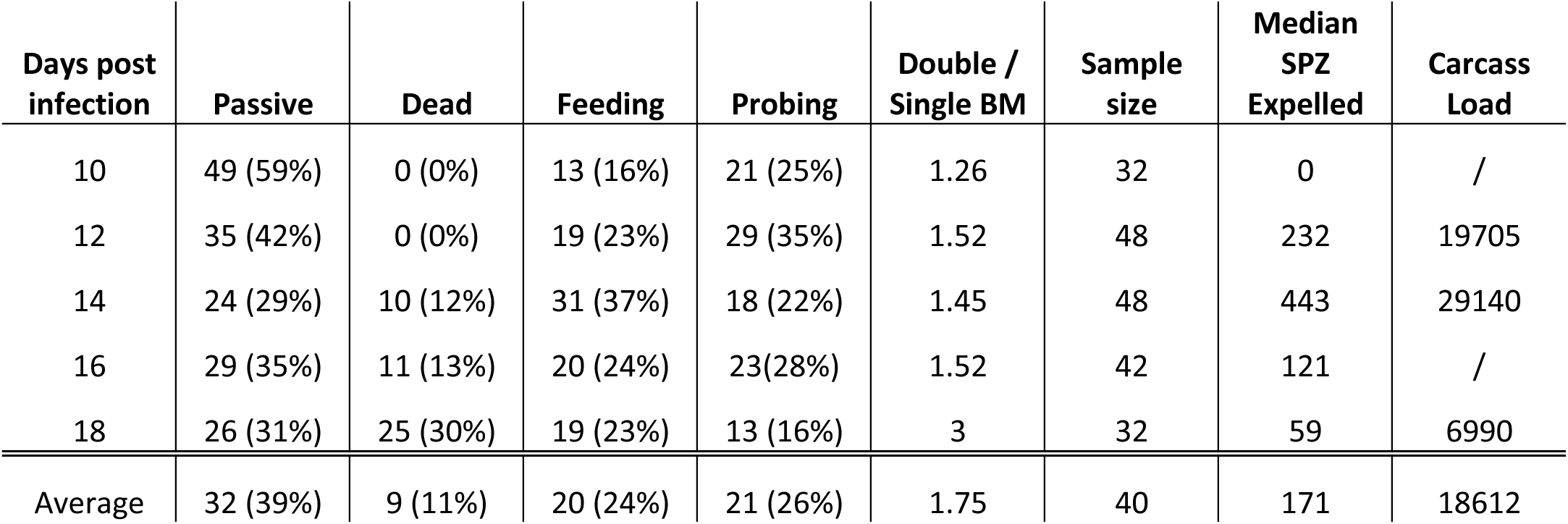
Continous timecourse experiment information Underlying mosquito cohort had an infection prevalence of 100% and an average 6.5 oocyst per midgut. Double/Single BM gives the ratio of interacting mosquitos that have received an additional uninfectious bloodmeal or not.

### A second bloodmeal at 6 dpi does not modulate inoculum size or timing

In natural settings, mosquitoes may take additional blood meals after their first infectious meal. Previous reports have suggested that a second bloodmeal, variously given from 3-9 days, after the initial infectious bloodmeal accelerates (but does not increase intensity of) parasite development^27, 28^ or synchronizes and thereby increases intensity of (but does not accelerate) parasite infection^29^, respectively. To explore if a second non-infectious bloodmeal impacts inoculum size or timing, mosquitoes were evenly split into only receiving the infectious bloodmeal or receiving an additional non-infectious bloodmeal at 6 dpi. In the discontinous variant we observed no statistically significant differences in inoculum size on any day when comparing the groups that received a single versus two blood meals (Fig. 4A). Notably, no evidence for earlier expelling was found and salivary gland loads were statistically indistinguishable when correcting for multiple testing. We also assessed timing of initial sporozoite colonization of salivary glands through microscopy and found no significant difference in the number of positive mosquitoes (Fig. 4B). For both salivary gland load and inoculum size, inclusion of a second bloodmeal in the GLM did not show alter the associations described above (Supp. Fig. 4). Contrary to our expecations, we observed lower infection intensity at 18 dpi for mosquitoes that received an extra bloodmeal (Fig. 4C). In conclusion, we find no evidence for a secondary bloodmeal influencing inoculum size or timing of expelling but some indications that a second blood meal lowers infection intensities starting at 18 dpi.

**Figure 4.**
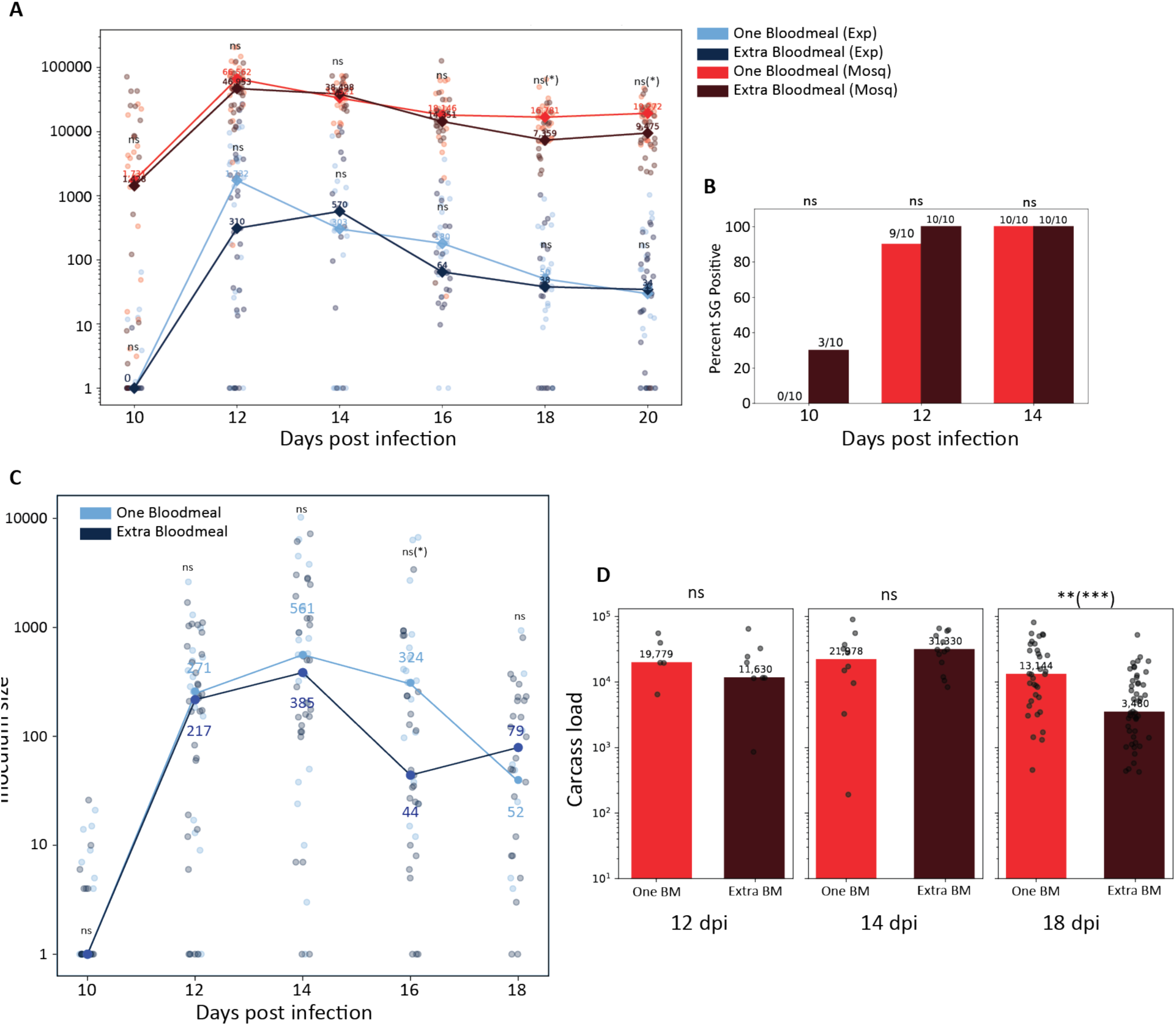
The effect of a second bloodmeal **A** Median salivary gland load and inoculum size from 10 dpi to 20 dpi for mosquitoes that have received the infectious bloodmeal only (light blue inoculum, light red salivary gland load) or an additional non- infectious bloodmeal (dark blue inoculum, dark red salivary gland load). An additional bloodmeal does not appear to significantly modulate inoculum size or SG load (two-sided MWU with Bonferroni correction) **B** Proportion of positive salivary glands in mosquitoes that have received one (light red) or an additional bloodmeal (dark red) on 10,12 and 14 dpi. No significant difference in salivary gland positivity between one and an additional bloodmeal was found (Fisher’s exact test with Bonferroni correction). **C** Relative frequency of different inoculum size ranges (0, 1-100, 101-500, 501-1500, >1500) and salivary gland loads (0, 1-5000, 5001-25000, 25001-100000, >100000) over time (10 – 20 dpi) in mosquitoes that did or did not received an additional bloodmeal. **D** Median inoculum size of mosquitoes from the continuous experiment that have received only the infectious (light blue) or an additional non-infectious bloodmeal (dark blue) **E** Carcass load of mosquitoes that died throughout the experiment (12 dpi, 14 dpi) or were collected at the end of the experiment (18 dpi) that received only the infectious bloodmeal (light red) or an additional non-infectious bloodmeal (dark red). On 18 dpi carcass load was significantly lower in mosquitoes that received an additional bloodmeal (two-sided MWU, *p* = 0.001). On 12 dpi and 14 dpi no differences were found (*p* = 0.0518, *p* = 0.402).

### Probing is sufficient for sporozoite expelling

The SpitGrid records the interaction of mosquitoes, allowing us directly examine the impact of different modest of interaction with the artificial bite substrate. We categorized the mosquitoes into “probing” (probing but no engorgement) and “feeding” (probing and engorgement) categories. We observed no consistent higher inoculum size for feeding compared to probing mosquitoes across the continuous and discontinuous experiments (Supp. Fig. 5A). Similarly, the frequency of non-expelling mosquitoes showed no evident difference between feeding or probing mosquitoes (Supp. Fig. 5B, C, D). Both feeding and probing-only mosquitoes followed the same general trajectory of peak expelling followed by a gradual decline across both experiments. To better account for the temporal nature of the data, we again applied a GLM approach on the pooled data of both continuous and discontinuous experiments and observed that the binary behavior classification is not a statistically significant predictor of inoculum size (Supp. Fig. 5D). We therefore conclude that mosquitoes that do not take a (full) bloodmeal can readily inoculate sporozoites and transmit malaria, consistent with previous findings^14^.

**Figure 5.**
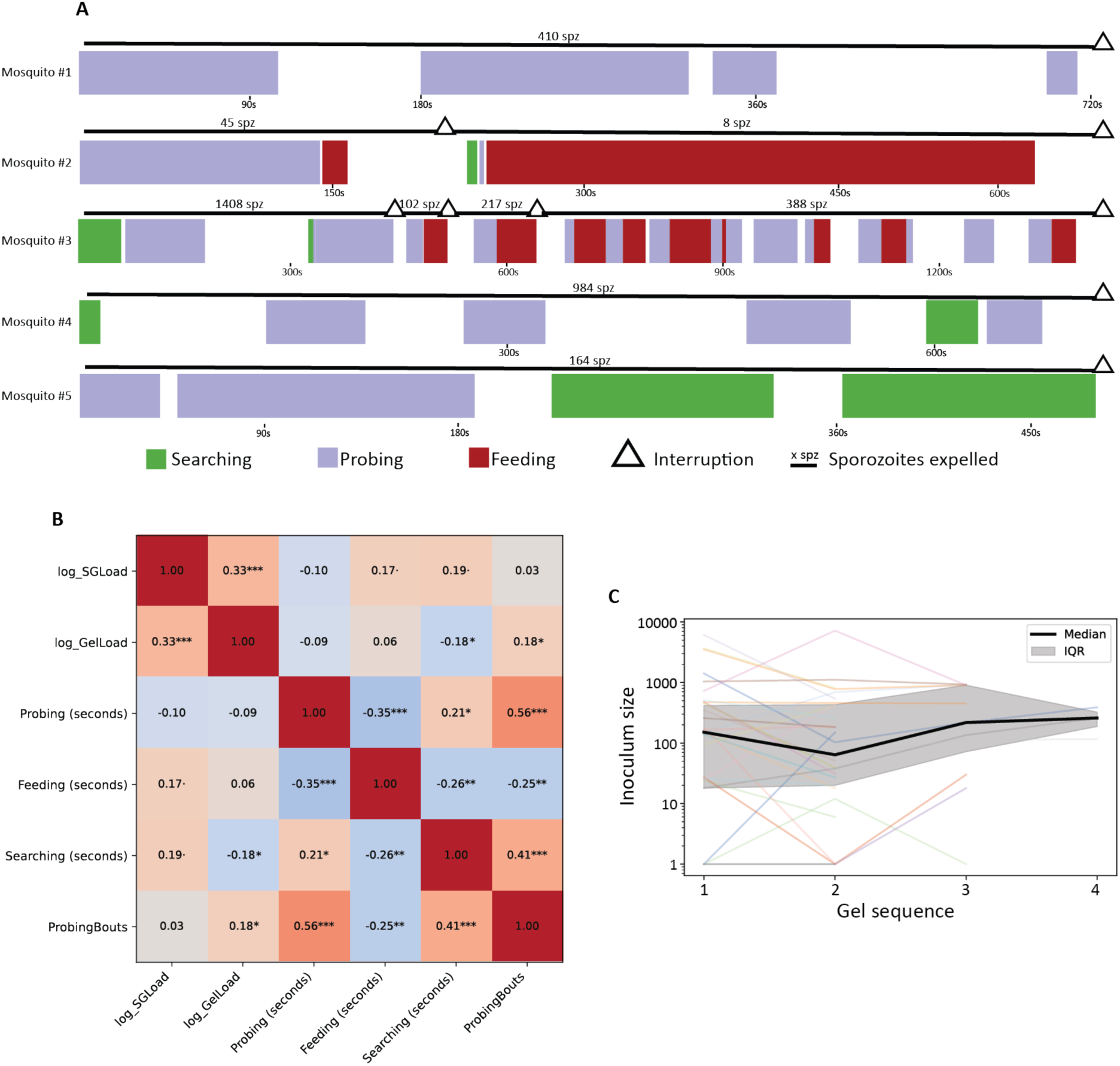
Behavioral determinants of sporozoite inoculum. **A** Representative ethograms from behavioral experiments. Each row shows one mosquito and the sequence and duration of behavioral states during a single exposure (Searching = green, Probing = purple, Feeding = red). Access to the artificial bite substrate was interrupted by lifting the cage off the gel (triangle), allowing the substrate to be exchanged. Sporozoite output quantified from each gel is annotated as the number of sporozoites expelled (x spz). **B** Correlation matrix of log-transformed salivary gland load (log_SGLoad), log-transformed gel load (log_GelLoad; inoculum size), probing time, feeding time, searching time, and number of probing bouts. Cells show correlation coefficients with significance indicated by asterisks. **C** Inoculum size across successive gels of interrupted interactions (gel order 1–4). Individual mosquito trajectories are shown as thin colored lines; the median (black) and interquartile range (grey) are overlaid, showing no systematic decline across consecutive interactions.

### Behavioral determinants of sporozoite inoculum

To investigate the behavioral determinants of incolum size in more detail, we modified the experimental setup to monitor a single mosquito in high resolution allowing quantitative assessment of bite-related behaviors. For each interaction, we quantified time spent interacting with the substrate without stylet insertion (searching time), time spent with stylet inside substrate but not engorging (probing time), count of initiation of probing behavior (probing bouts), and time spent engorging (feeding time). To better link inoculum size and observed behaviors, we interrupted mosquito-substrate interactions at arbitrary points, by lifting the cage off the artificial bite substrate. A new artificial bite substrate was provided and the mosquito could resume its interaction. These interruptions further allowed us to interrogate whether repeated probing or feeding in a short time window reduce the inoculum size of subsequent bites. For each mosquito, salivary gland load was determined alongside sporozoite quantities in the artificial bite substrates. Interruption experiments were performed on 83 individual mosquitoes including three different mosquito species (Supplementary Information 1). To illustrate the structure and diversity of mosquito–substrate interactions, we visualized behavioral sequences as ethograms showing behavioral states (searching, probing, feeding), interruptions, and corresponding sporozoite outputs per interaction. Ethograms of 5 reperesentative mosquitoes (Fig. 5A) highlight the heterogeneity in behavior across mosquitoes, indicate the absence of a clear depletion trend across successive gels, and the fact that sporozoite deposition can occur both with and without engorgement.

Overall, no single parameter alone explained much of the variability in inocolum size, pointing to a multifactorial process (and likely a large stochastic component) (Fig. 5B). Of the behavioral variables, more probing bouts resulted in significantly larger inocula, while longer searching time was significantly associated with smaller inocula. Surprisingly, probing or feeding time did not emerge as a predictive variable. Saliavary gland load remained the strongest predictor of inoculum size, which is in agreement with previous literature and our timecourse data (Supp Fig. 2A, Fig. 2E).

When comparing mosquito substrate interactions that contained feeding to those that only contained probing, there was no difference in inoculum size (Supp. Fig 6A), in line with the timecourse data. Similarly, mosquitoes that did end up feeding in any of their interactions did not expell significantly more than mosquitoes that only probed (Supp. Fig 6B). Inoculum sizes of subsequent interactions were correlated (Supp. Fig. 6C) and median inoculum sizes did not decrease in later interactions compared to earlier interactions (Fig. 5C, Table 3). Suggesting that, also when periods between bites are much shorter there is no evidence of depletion, mirroring the timecourse data.

**Figure 6.**
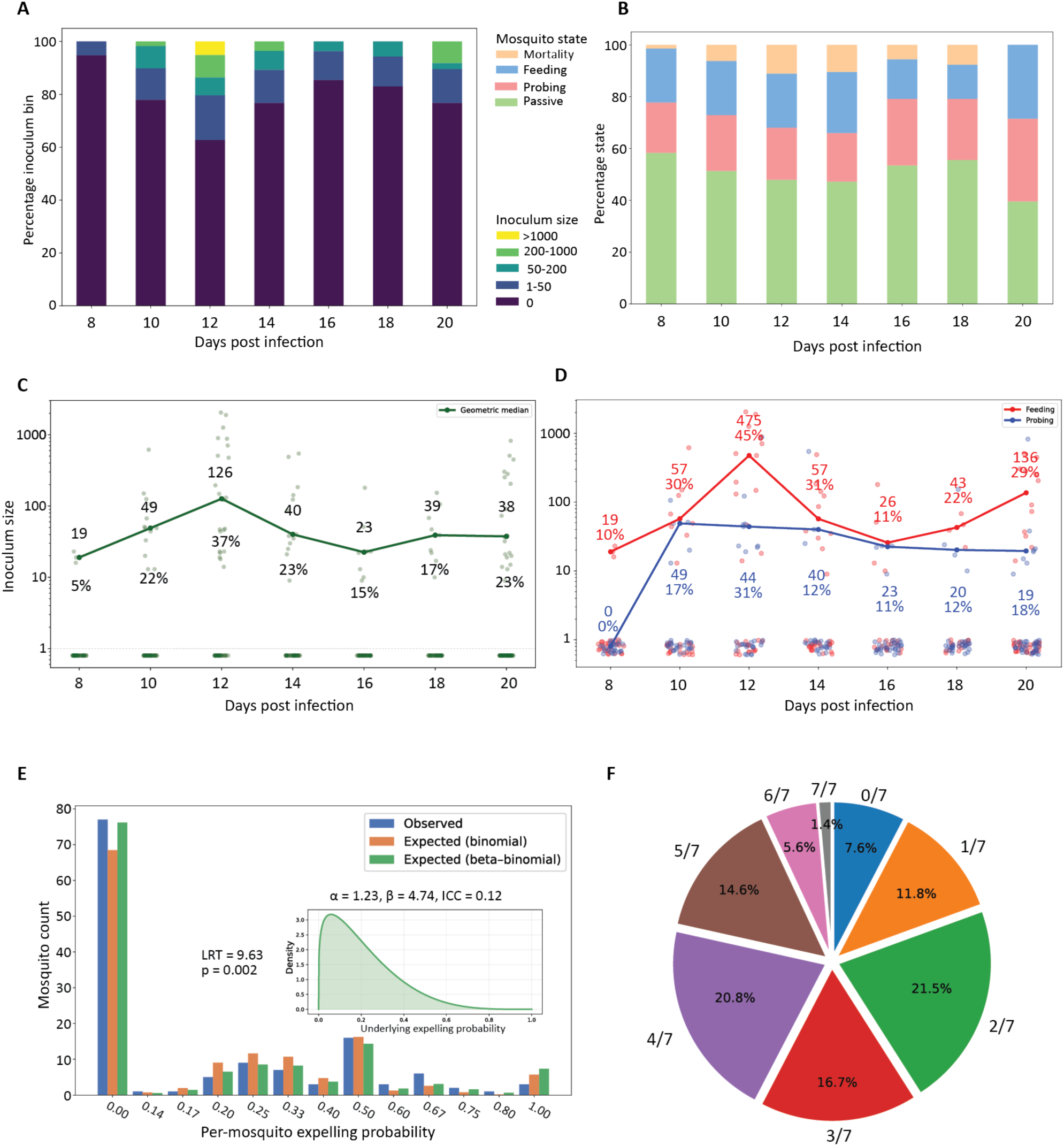
Inoculum dynamics of *Plasmodium vivax*. **A** Relative frequency of inoculum size ranges over time (8–20 dpi; 0, 1–50, 50–200, 200–1000, >1000 sporozoites). **B** Relative frequency of behavioral outcomes during each exposure (Passive, Probing, Feeding, Mortality). **C** Inoculum size per mosquito per day (log scale; individual datapoints as transparent circles) with the geometric median per day (green line; values annotated) and the daily expelling rate (percentage of mosquitoes with detectable sporozoites in the bite substrate; annotated below each day). **D** Same as (C), stratified by behavior (Feeding, red; Probing, blue); geometric medians and behavior-specific expelling rates are annotated. **E** Observed distribution of per-mosquito expelling frequency across the 7 exposure days compared to expectations under a binomial model (shared expelling probability) and a beta-binomial model (mosquito-to-mosquito variability in expelling probability); inset shows the fitted beta distribution (α, β) and inferred intraclass correlation coefficient. **F** Pie chart showing the fraction of mosquitoes expelling on n of 7 exposure days (0/7 to 7/7).

**Table 3.**
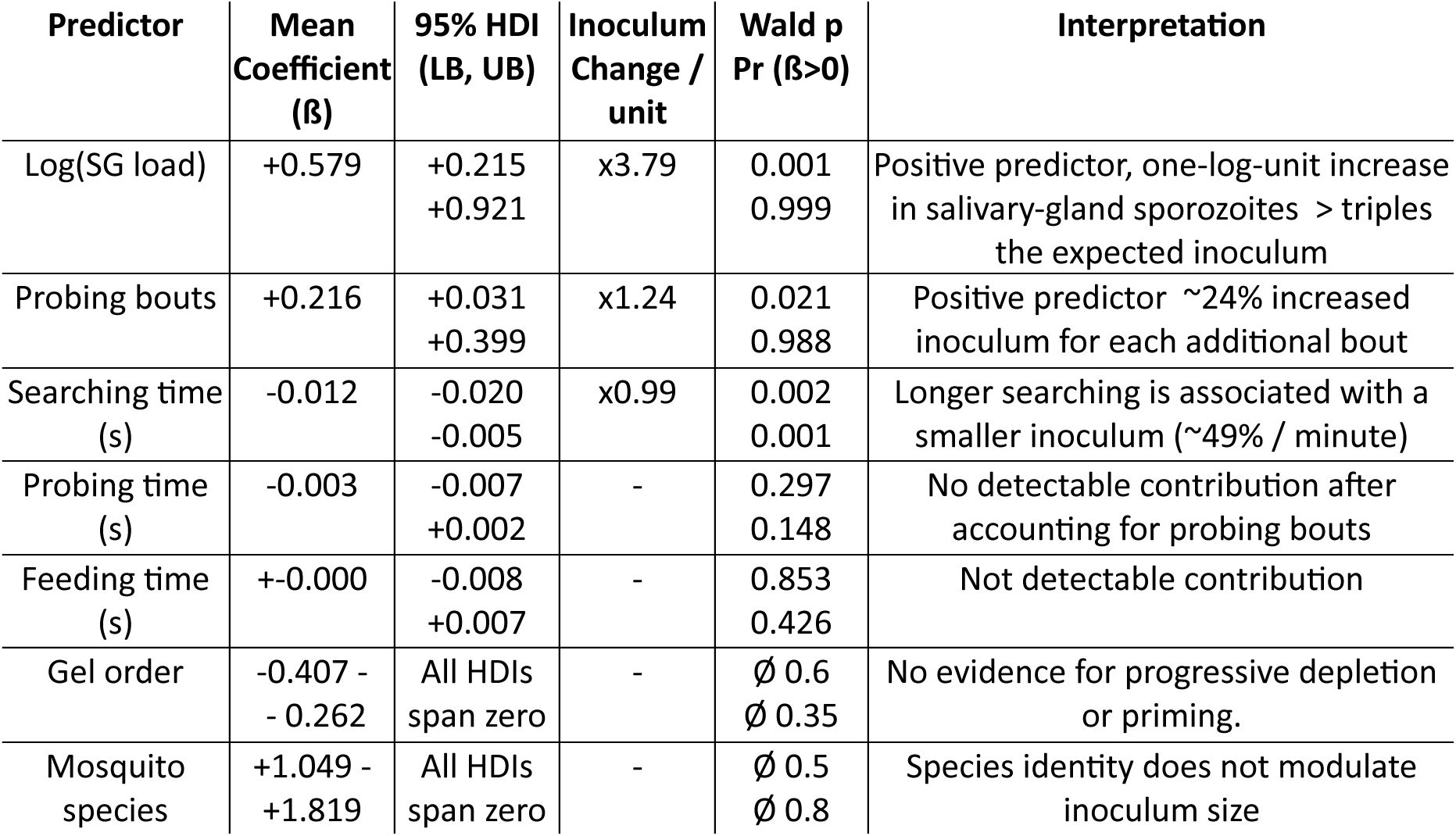
Gaussian Mixed effect model to approximate determinant of incolum size for P. falciparum. Coefficients (β) are shown with 95% highest density intervals (HDI), corresponding fold-change in expected inoculum per unit increase of the predictor, and Wald test statistics. Pr(β > 0) is the posterior probability that the regression coefficient is positive. Positive effects indicate larger inocula, negative effects indicate reduced inocula

As single variables appear to only explain a small part of the observed variation we next employed a Gaussian mixed effect model to account for the multifactorial nature of sporozoite delivery. We implemented the model with fixed predictors that contained all variables with a plausible mechanistic link to inoculum size (salivary-gland load, probing bouts, feeding time, probing time, searching time, mosquito species, bite substrate order) as well as random intercepts to account for experiment day variation (Table 3). Through this analysis we found that salivary-gland load remained the strongest predictor with every 10- fold increase being associated with a ≈3.8× higher inoculum. Number of probing bouts showed a positive association with a 24% larger inoculum for each additional probing bout. Searching time also remained a negative predictor with the inoculum halving for each minute that the mosquito spent searching. Interestingly, both feeding and probing time had no measurable impact on the inocolum size. The fixed effects together explained ≈29% of variance and adding the day random intercept raised this to 48%. The remaining variation may be caused by anatomical variation between individual mosquitoes, qualitative differences in probing or feeding behavior, or aspects of the infection not captured by salivary gland load.

### *P. vivax* Inoculum sizes are smaller than *P. falciparum* but show similar temporal dynamics

Despite its large medical significance, basic properties of the infectious bite such as inoculum size and the temporal onset of transmissibility remain unaddressed for *P. vivax*. This is largely due to the fact that this species cannot be maintained in continuous culture, and the resulting scarcity of experimental infections. To close this knowledge gap, we leveraged a clinical trial in which human participants were experimentally infected with *P. vivax* with the aim to infect *An. stephensi* mosquitoes. We took advantage of this unique sitiuation to measure inoculum size, kinetics and behavioral determinants of *P. vivax* transmission. Based on limited historical data, the EIP of *P. vivax* has been suggested to be around two days shorter compared to *P. falciparum*^30, 31^. We therefore characterized sporozoite expelling from 8-20 dpi using a continous time course. To limit mortality in the context of the limited supply of *P. vivax* infected mosquitoes, mosquitoes were not starved prior to artificial bite substrate exposure. Indeed, mortality remained below 10% throughout the experiment (Fig. 6B). Compared to *P. falciparum*, *P. vivax*-infected mosquitoes expelled markedly smaller inocula and a substantially larger proportion of mosquitoes showed no detectable sporozoite expulsion (Fig. 6A, C). To accomodate the larger fraction of mosquitoes not expelling detectable sporozoites, we analyzed geometric medians (excluding zeros due to log transformation) and frequency of expelling as summary metrics of expelling activity over time. *P. vivax* expelling followed the same general trend as observed for *P. falciparum* showing an early rise in inoculum size, peaking at 12 dpi, and a post-peak decline. Relative to *P. falciparum,* temporal dynamics of *P. vivax* expelling seemed less pronounced, with median inoculum sizes and expelling frequencies varying approximately five (compared to ∼30-fold in *Pf*) and two-fold, respectively (Fig. 6A). The reportedly shorter EIP of *P. vivax* was reflected in comparitively large inocula and expelling frequencies at 10 dpi. While we did find detectable inocula already at 8 dpi in three mosquitoes, they were small and represent only 5% of the interacting population, suggesting limited transmission relevance at this early timepoint. While carcass load at 20 dpi did not seem to correlate with inoculum size or expelling frequency (Supp. Fig. 7C,D), mosquitoes originating from cohorts with a higher mean oocyst load appeared to have a consistently higher expelling prevalence but no consistent effect on inoculum size (Supp. Fig. 7E,F). Next, we investigated whether expelling probability was uniformly distributed across mosquitoes, or if some individuals were inherently more likely to expel than others. Analyzing subsequent interactions, we found that mosquitoes that previously interacted without expelling, were around half as likely to expel during the following interaction compared to those that did expel previously, suggesting differences in expelling between mosquitoes (Supp. Fig. 8A, B). To further test for individual heterogeneity in expelling frequency, we compared a simple binomial model, where all mosquitoes expel with the same likelihood, to a beta–binomial model that includes an overdispersion parameter, allowing for differences in expelling frequency between individuals (Fig. 6E). The overdispersion model is a significantly better fit to the observed data, suggesting heterogeneity between mosquitoes in the likelihood of expelling. In line with this, when plotting trajectories of individual mosquitos consistent non-expellers are more likely in frequently interacting individuals than one would expect and we frequently observe repeated expelling by individuals (Supp. Fig. 8C, D). We furthermore observed an effect of feeding behavior on *P. vivax* expelling. Across all days, mosquitoes that engorged showed a higher likelihood of expelling and larger median inocula compared to mosquitoes that only probed (Fig. 6D). Combining all plausible modulators of inoculum size or expelling frequency in a GLM, corroborated findings from separate analysis (Table 5, Table 6). Taken together, the *P. vivax* data suggest a quantitatively different transmission profile compared to *P. falciparum* showing less pronounced temporal dynamics of expelling, smaller inocula, and a larger proportion of non-expellers. *P. vivax* showed a more pronounced influence of engorgement on both inoculum size and expelling frequency.

**Table 5.** GLM to approximate determinants of incolum size for P. vivax Coefficients (β) are shown with 95% highest density intervals (HDI), corresponding fold-change in expected inoculum per unit increase of the predictor, and Wald test statistics. Pr(β > 0) is the posterior probability that the regression coefficient is positive. Positive effects indicate larger inocula, negative effects indicate reduced inocula

| Predictor | Mean Coefficient ($\beta$) | 95% HDI (LB, UB) | Inoculum Change / unit | Wald p Pr ($\beta > 0$) | Interpretation |
| --- | --- | --- | --- | --- | --- |
| Time pre 12 dpi | +0.954 | +0.857<br>+1.051 | x2.60 | <0.001<br>>0.999 | From 8-12 dpi mean inoculum increases with a factor of 2.6 per day |
| Time post 12 dpi | -0.234 | -0.369<br>-0.097 | x0.79 | <0.001<br><0.001 | From 12-20 dpi inoculum size decreases by around 20% per day |
| Feeding | +1.599 | +0.824<br>+2.375 | x4.95 | <0.001<br>>0.999 | If feeding around five times more sporozoites are expelled |
| Mean oocyst | +0.137 | +0.011<br>+0.263 | x1.15 | 0.04<br>0.984 | For every additional oocyst in cohort 15% larger inoculum |
| Carcass Load | +0.001 | +0.016<br><+0.001 | x1.00001 | 0.001<br>>0.999 | Every 100 additional parasites in carcass at 20 dpi, 1% larger inoculum |

**Table 6.**
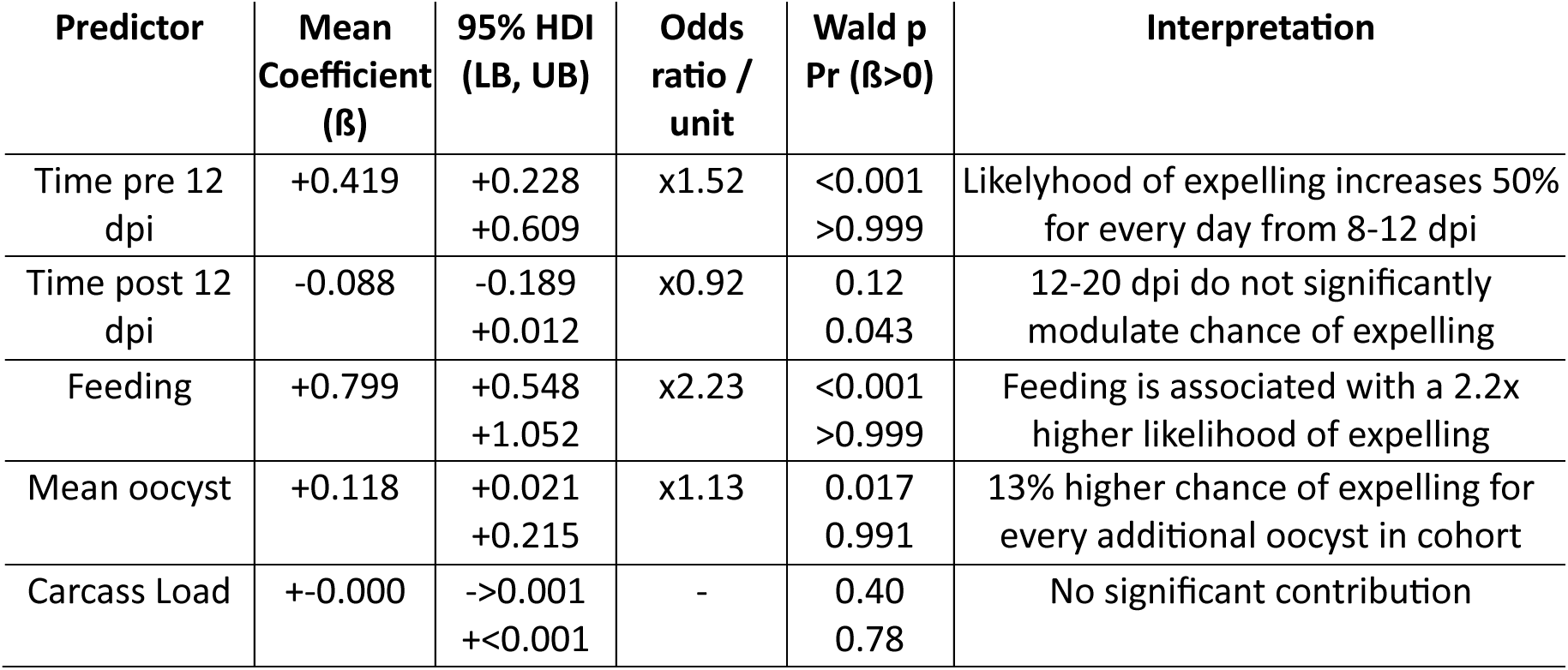
GLM to approximate determinants of expelling likelihood for P. vivax. Coefficients (β) are shown with 95% highest density intervals (HDI), corresponding odds ratios per unit increase of the predictor, and Wald test statistics. Pr(β > 0) is the posterior probability that the regression coefficient is positive. Positive effects indicate a higher likelihood of expelling, negative effects indicate a reduced likelihood.

## Discussion

We demonstrated the SpitGrid, a novel experimental platform enabling non-destructive high-throughput quantification of sporozoite expelling by individual mosquitoes over time, while capturing metrics related to feeding behavior. Using this platform, we characterized expelling dynamics of mosquitoes infected with the most important *Plasmodium* species, *P. falciparum* and *P. vivax*. Our data reveal that inoculum size is highly dynamic, peaking shortly after the extrinsic incubation period (EIP) has elapsed. Interestingly, for both parasite species we observed a significant decline in inoculum size post peak. We observe that probing alone, without engorgement, is sufficient to deliver sizeable inocula, and repeated feeding or probing at shorter or longer intervals, does not measurably reduce inoculum size. We demonstrate that engaging in probing more often increases inoculum size, whereas probing time or whether it results in engorgement did not significantly impact inoculum size. Together, these findings challenge the assumption that per-bite transmission potential is constant after the EIP has elapsed, and provide a nuanced understanding of physiological, temporal, and behavioral factors determining inoculum size. These novel temporal dynamics have immediate implications for how mosquito infectiousness is represented in epidemiological models and for how transmission-blocking interventions are evaluated.

Current transmission models largely treat infected mosquitoes as binary transmitters. Once the extrinsic incubation period has elapsed, each bite is assumed to carry a fixed transmission probability until the mosquito dies^32, 33^. We, however, observe clear temporal dynamics, which are likely highly relevant for the likelihood of establishing infection. In rodent models, mosquitoes with a higher salivary gland load are more likely to establish a successful infection^14^. In humans, the history of malariotherapy and controlled human malaria infection studies are also indicative of a positive correlation of dose with disease severity, shorter prepatent periods, and ability to break through protective immunity ^2, 34^. While the exact relationship remains poorly defined, sporozoite inoculum is evidently highly relevant for malaria transmission.

By directly measuring inoculum over time in individual mosquitoes, we show that infectious capacity has a strong temporal component. Sporozoite expelling peaks in a narrow window and then declines approximately log-linearly. If per-bite transmission probability roughly follows this inoculum profile, then models that assume lifelong, constant infectiousness will overestimate the contribution of the long mosquito survival tail to onward transmission^35^. These temporal dynamics also matter directly for how we evaluate interventions acting on parasite development in the mosquito. Tools that lengthen the extrinsic incubation period, such as gene drives that retard parasite development^36, 37^, temperature effects^38^, or certain antimalarials^39, 40^ and Wolbachia-based approaches^41^, are typically judged by the fraction of mosquitoes surviving long enough for onward transmission. If infectiousness is concentrated in a short, early high-inoculum window, it is unclear if a delay in parasite development changes how many bites fall in that window. The SpitGrid provides one side of this relationship by making inoculum directly measurable at scale.

### Revisiting the relationship between infection intensity and inoculum size

The divergence of inoculum size and salivary gland load after the initial peak suggests that supply, on the whole salivary gland level, and output are at least partially decoupled and become more so over time (Fig. 2A, F, Supp. Fig. 2A). Prior work has shown a rapid loss of sporozoite motility and infectivity after isolation from salivary glands^42^.^42^ This putative decay in function or aging of sporozoites could underlie a lower fraction of total salivary gland parasites that can be mobilized for expelling as previous work has shown that sporozoites need to actively move from the salivary glands to the ducts, to become available for expelling^46^.

Sporozoites have been described in various assemblies inside the salivary glands, from large aligned clusters, to more disorganized assemblies to small groupings or individual sporozoites^43–45^. If sporozoites aggregate progressively over time, which may reduce their ability to move towards the salivary duct, the measured salivary gland load could overestimate the expellable pool as time progresses. Investigating whether there are shifts in sporozoite assemblies which could influence expelling dynamics is an interesting avenue to mechanistically understand sporozoite expelling over time.

Salivary glands themselves have also been shown to experience variable disruptions from sporozoite invasion. While some salivary glands remain intact, others show structural perturbations and depletion of salivary proteins^45^. Recent work has shown extensive tissue damage at 19-21 dpi that is characterized by ruptured barriers, local inflammation, and leakage of salivary proteins into hemolymph^47^. These perturbations could cause defects in salivation and consequently negatively affect sporozoite expelling. Exploring this hypothesis will require integration of salivation volume measurements into the SpitGrid and assessments of infected salivary gland anatomy across different timepoints and infection intensities.

### Potential underpinnings of heterogeneous inoculum size

On any given day post EIP we, and previous works^12, 13^, observe a large heterogeneity in inoculum size or even lack of any expelling among mosquitoes with similar infection intensity and feeding behavior. Initial observations of this apparent stochasticity in inoculum size have linked it to clustering of sporozoites and suggested it as a strategy of the parasites to pass the minimally infective dose^48^. Our repeated measurements on individual mosquitoes within minute-scale timeframes show a high degree of correlation and lower variability within an individual mosquito (Supp. Fig 6C), suggesting that features of an individual mosquito or specific infection trajectory influence inoculum size. Recent imaging of infected salivary glands suggests plausible tissue-level explanations for this heterogeneity^45^. Even at similar parasite burdens, salivary glands can range from structurally intact to markedly perturbed, with variations in duct or lumen and infection-associated damage that could either constrain sporozoite release (yielding non-expellers) or, conversely, increase sporozoite access to secreted saliva (yielding large inocula). Consistent with this, modeled compared to observed values are least accurate at zero-inoculum and very high inoculum events (Supp. Fig. 9A). Mirroring this tissue level and inoculum heterogeneity, measurements of salivation volume, also find up to 400 fold differences between individual mosquitoes, when salivating into mineral oil^6^. Together, these observations suggest that mosquito-to-mosquito differences in salivary-gland anatomy, infection-associated tissue disruption, and salivation heterogeneity may impose strong, individual constraints on sporozoite release that are independent of specific parasite quantities.

### On the development of inoculum size over time

Our time resolved and repeated measurements from individual mosquitoes reveal that sporozoite delivery peaks after the onset of infectivity and then declines over time at the population level and across successive bites by the same mosquito. To our knowledge, these data constitute the first longitudinal, single-mosquito description of inoculum dynamics.

Although prior work did not quantify inoculum size directly or follow indivuals over time, historical observations provide interesting parallels. Barber and colleagues reported “degenerate” sporozoites accumulating in older infections, with degeneration appearing earlier and progressing more strongly under warmer conditions, and noted that the presence of sporozoites in salivary glands is not necessarily a reliable criterion for infectiousness^49^. Other reports linked older infections and degenerate *P. vivax* sporozoites to reduced infectivity in human exposures ^19^. These observations are consistent with the idea that infectiousness can decay even when salivary glands remain sporozoite-positive. Mechanistically, our data are compatible with at least two non-exclusive scenarios. First, as discussed above^42^, parasites may age within the salivary glands, progressively losing fitness such that a smaller fraction can enter the ducts and become available for expelling. This would naturally produce a decline in inoculum size even if total salivary-gland load decreases only modestly. Second, the mosquito may increasingly limit or kill parasites over time e.g. through local immune activity in salivary tissues^50^ or the aforementioned infection-associated tissue damage, constraining expelling without necessarily being reflected in salivary gland-associated parasite DNA. Temperature could plausibly modulate both mechanisms. If decay is dominated by parasite aging, higher temperatures may accelerate metabolic turnover and functional decline. If decline reflects mosquito-mediated factors, temperature could modulate host defences or exacerbate salivary-gland pathology, again producing earlier loss of effective infectiousness. In line with this, Shapiro and colleagues observed a temperature-dependent rise and fall in the fraction of salivary-gland-positive mosquitoes at ≥27 °C, with higher temperatures producing an earlier and more compressed trajectory^38^. While their readout was infection prevalence (not inoculum size), it is indicative of temporal patterns in infectivity and temperature as an important modulator. Distinguishing these scenarios will require integrating inoculum measurements with markers of sporozoite viability and mobility, and longitudinal assessment of salivary-gland anatomy and salivation output across temperatures and infection intensities.

### Caveats of extrapolating *P. berghei* transmission data to human malaria

Our understanding of sporozoite biology relies heavily on the rodent parasite *Plasmodium berghei*, yet several pieces of evidence, as has been suggested by Vezier et al.^51^, indicate that its sporozoites are different from those of *P. falciparum* in ways that directly affect temporal patterns of infectiousness. *P. berghei* sporozoites retain high infectivity for extended periods^51, 52^. Single-cell transcriptomic analyses reinforce this stability, showing that *P. berghei* sporozoites enter a quiescent state within the salivary gland mounting a translational and metabolic upshift only upon being salivated^53^. In *P. falciparum*, by contrast, transcriptomic differences between salivary-gland and salivated sporozoites are modest^54^, and recent work on sporozoite ageing show a rapid decay in motility and infectivity^42^. A parsimonious interpretation is that *P. berghei* sporozoites conform to a quiescent salivary-gland state optimized for longevity, while *P. falciparum* sporozoites remain constitutively primed for host invasion, possibly sacrificing long-term stability.

### How different are transmission dynamics of *P. vivax* and *P. falciparum*?

Compared with *P. falciparum*, far fewer data exist on mosquito-to-human transmission for *P. vivax*. Controlled human malaria infection (CHMI) studies typically expose volunteers to 3–5 *P. vivax*–infected mosquitoes (same number as *Pf* designs^55^), a number chosen to reliably induce blood-stage infection rather than to probe dose–response behaviour^56^. A systematic re-analysis of historical malariotherapy experiments found limited association between the number of infected bites and the prepatent period, whereas parasite strain and vector species were much stronger determinants^57^. Field work from Papua New Guinea further shows that, at similar entomological inoculation rates for *P. falciparum* and *P. vivax*, *P. falciparum* infections are substantially more prevalent^58^. Burkot and colleagues concluded that *P. falciparum* is more efficiently transmitted from mosquito to human and suggested higher sporozoite quantities in wild-caught *P. falciparum*–infected mosquitoes as a likely contributing factor. They further suggest that *P. vivax* produces 10 fold less sporozoites per oocyst but the only other paired dataset suggests no meaningful difference between the two species while recent investigations on *P. falciparum* suggest more sporozoites per oocyst than suggested by either comparative study^12, 13, 58, 59^. In this context, our *P. vivax* SpitGrid experiment, showing smaller inocula, a higher proportion of non- expelling bites and markedly lower carcass parasite loads than in our *P. falciparum* infections (Supp. Fig. 9B) despite comparable mean oocyst quantities, would be consistent with Burkot et al.^58^. To what extent these findings are compatible with similar performance of both species in CHMI trials or whether other mechanisms such as more efficient skin traversal or liver infection by *P. vivax* might be at play, remains to be seen. A future implementation of the SpitGrid to match specific salivary gland loads to inocula for *P. vivax* or explore the impact of vector species , can help us tease apart how much of our findings are based on intrinsic differences between biology of the two species and how much is owed to the specific circumstances in this study.

### Behavior, depletion, and inoculum

Transmission requires both the presence of sporozoites in the salivary glands and a biting interaction with the host, and it is intuitive to assume that the mechanics of the latter influence the dose delivered. A second intuition is that repeated delivery events within a relatively short duration, which are commonplace in nature^60^, may progressively deplete the expellable pool. Prior studies, working with limited numbers of mosquitoes, have challenged these assumptions. Interrupting a bite and allowing mosquitoes to resume on a new mouse skin does not reduce subsequent sporozoite output^8^ and probing on a mouse before forced salivation does not diminish the number of sporozoites expelled into mineral oil^61^, and with *P. berghei* parasites probing alone can establish infection as effectively as full engorgement^62^. Our data provide a broader and time-resolved test of these ideas. Counterintuitively, neither feeding time nor probing time was strongly associated with inoculum size, whereas the number of probing bouts showed the clearest positive association. This fits with an earlier observation that sporozoites are disproportionately expelled at the onset of salivation^6^ suggesting that once probing is initiated, most of the inoculum is delivered immediately, contrary to earlier models that assumed a linear, steady outflow of parasites^63^. Consistent with prior experimental work, we also find no evidence for meaningful depletion^8^. Repeated probing or feeding bouts, whether minutes apart or across multi-day intervals, do not reduce subsequent inoculum size. Taken together, these results indicate that the behavioral determinants of expelling are governed more by the initiation of probing than by feeding duration and that the expellable pool is remarkably robust to repeated biting opportunities. It would be highly informative to test how these observations interact with the behavioral alterations induced by insecticide resistant genotypes and insecticide treated bednets^15–17^.

### Validation, methodological advances and outlook

The SpitGrid overcomes major limitations of existing approaches to quantify sporozoite delivery. Classical forced-salivation methods are a poor proxy for natural biting behavior, produce unnaturally low inocula in *Plasmodium*^7, 9, 10^, and are not predictive of infection outcomes in arboviruses^64^. Skin-based assays yield realistic doses but are labor-intensive, low-throughput, or rely on animals^12, 13, 63^. Their opaque nature also makes nuanced behavioral analysis challenging.

The SpitGrid adresses these issues, yet an important question is whether the artificial bite substrate provides a realistic proxy for *in vivo* expelling. Andolina *et al.*^13^ report median inocula of 150 – 1000 sporozoites using blood-soaked dermal matrices and 11 % - 47% of non-expelling but infected mosquitoes, and Kanatani *et al.*^12^ find a geometric mean of 850 sporozoites among transmitting mosquitoes and ∼14% non-transmitters using mouse skin excision. Our agarose-based system yielded median inocula of 200-1000 sporozoites and a 15% rate of non-expellers in the same period post infection. These comparisons indicate that our measurements fall within the physiological range observed in true skin-based assays and support the SpitGrid as a realistic proxy for *in vivo* expelling.

By coupling a simple transparent gel substrate with physiologically relevant cues to a modular housing array the SpitGrid is inexpensive, scalable, and compatible with imaging and repeated measures on the same mosquito. The single-mosquito variant further allows high-resolution behavioral readouts and mouthpart-level analysis, enabling linking of fine-grained behaviors to sporozoite expelling. As mosquitoes can be maintained in the grid and repeatedly challenged, the platform naturally accommodates before-and-after designs, for example the addition of sugar-borne interventions. Its modular structure makes it readily adaptable for testing transmission-blocking tools that alter parasite development rather than parasite numbers^65^ and for examining behavioral shifts induced by aforementioned interventions^15–17^. Finally, the same principles can likely be extended to quantify expelling of mosquito-borne viruses or to other pathogen-transmitting insects, making the SpitGrid a broadly applicable platform for vector– pathogen research. By making the inoculum itself a directly measurable trait at scale, the SpitGrid resolves a long-standing blind spot in transmission biology.

## Acknowledgements

We thank the members of the Vector Biology team for valuable input on data analysis and experimental design. We thank Jolanda Klaassen, Laura Pelser-Posthumus, Astrid Pouwelsen and Saskia Mulder for breeding of mosquitoes and handling of the infected mosquitoes. We thank Rianna Storter for culturing parasites used for mosquito infection and Karina Teelen for DNA isolations and qPCR work. This works was supported by the Netherlands Organisation for Scientific Research (NWO-VI.Vidi.213.167) and an Hypatia fellowship to FJHH, the European Research Council (ERC-CoG 864180, QUANTUM) to TB. *P. vivax* experiments were supported by the OptiViVax project (Horizon Europe 101080744) and co-funding from UK Research and Innovation (UKRI) under the UK government’s Horizon Europe funding guarantee and Swiss Government’s State Secretariat for Education, Research, and Innovation (SERI).

## Material & Methods

### Mosquito rearing

*An. stephensi* (Nijmegen Sind-Kasur strain)^66^ mosquitoes were maintained at 26°C and 80% relative humidity under a 12 h:12 h reverse light-dark cycle, with ad libitum access to 10% (w/v) glucose provided on soaked cotton pads. For the behavioral experiments *An. coluzzii* (N’gousso strain)^67^, *An. gambiae* s.s (Kisumu strain)^68^ mosquitoes were reared in the same conditions.

### *Plasmodium falciparum* culture & mosquito infection

NF54 *P. falciparum* gametocytes were generated in an automated shaker system as previously described^69^. On day 14 after gametocyte induction, cultures were fed to 1–5-day-old female *An. stephensi* using a midi- feeder setup as previously described^70^. Prior to start of SpitGrid experiments, infection prevalence and average oocyst intensity were assessed by dissecting midguts from 10 mosquitoes. Unless stated otherwise, mosquitoes received a second, non-infectious bloodmeal 6 days after the infectious bloodmeal.

### *Plasmodium vivax* mosquito infection

*P. vivax* strain PvW1 (Thailand) cryopreserved stabilates of asexual blood stages^71^ were used to infected malaria-naïve participants (Study ID NL-OMON57011 for more details). Between day 11 to day 25 after inoculation, blood was sampled from parasitemic participants that were detected to be parasitemic and was used to feed *An. stephensi* mosquitoes, using the procedures described for *P. falciparum* infections above.

### Fabrication of SpitGrid components

All SpitGrid components were constructed using a laser cutter (Trotec Q400, CO_2_ laser). Vector design files for recreating the SpitGrid are available via Figshare (10.6084/m9.figshare.31007197). The cage-grids (4×6 array) were constructed from 3 mm transparent acrylic (VikuGlass XT), with a perforated 1.5 mm acrylic bottom to allow mosquito access to the artificial bite substrate. The top openings were covered with a double layer of slitted dental dam (HySolate Fiesta, Colgene), with the slit orientations of the two layers rotated by 90°, to prevent escape when mosquitos are added to the SpitGrid. The dental dam was laser-cut and replaced between experiments. For the bite-substrate grid, a matching pattern was cut from 1.5 mm acrylic and mounted on a 5 mm acrylic carrier plate using a thin layer of polydimethylsiloxane (PDMS, SYLGARD 184, cured at 65°C for 2 h). To maintain skin-like substrate temperature (35°C), indium tin oxide (ITO)-coated glass (Saint-Gobain) was used as a transparent heater controlled by a temperature controller with an external probe. The heated glass was mounted on optical rails and a Basler a2A2840-48umPRO camera equipped with a 8 mm lens (Navitar F1.4) was positioned beneath for behavioral imaging. Homogenous illumination was provided with an LED panel mounted above the SpitGrid (Godox LEDP260C).

For behavioral experiments, a modified setup with a single small cage, made from 3 mm transparent acrylic with a perforated 1.5mm acrylic bottom (VikuGlass XT), was used (Figshare stable link). For higher resolution filming a Basler acA4024-29uc camera with a 100 mm lens (Canon macro EF 100 mm f/2.8L) was mounted beneath. The artificial bite substrate was cast into a small PDMS container. A smaller transparent heater, with the heated area being the size of the bite substrate, was custom made by patterning ITO- Coated PET film (Thorlabs OCF2520), with a laser cutter and, together with copper wires, sealing it with adhesive acrylic film (Thorlabs OCA8146-2). The wires were then connected to a power supply (Instek GPD- 3303D) and voltage for the target temperature was empirically determined and confirmed with a thermal camera (FLIR C5).

### Generation of artificial bite substrates

To minimize ATP hydrolysis or other deterioration over time, grids with artificial bite substrates were prepared the morning of each experiment. A bulk agarose solution was prepared to final concentrations of 150 mM NaCl (Sigma-Aldrich, S7653), 25 mM sodium bicarbonate (Sigma-Aldrich, 31437-M), 4.5 mM glucose, and 0.6% (w/v) UltraPure LMP agarose (Invitrogen #16520-050) and pH was adjusted to 7.4. Components (except ATP) were heated until the agarose fully dissolved and the solution was allowed to cool to approximately 35°C before ATP addition (Sigma-Aldrich, A3377, 1 mM final concentration). The gel solution was then cast into the bite-substrate grid and covered with a flat glass/acrylic plate to ensure a smooth surface and consistent volume across cells. Grids were incubated at 4°C for 30 min to ensure complete gelation.

### SpitGrid assays and experimental designs

SpitGrid experiments were conducted at 21°C and 80% relative humidity; outside assay periods, mosquitoes were maintained under standard insectary conditions (27°C, 80% RH, 12 h:12 h cycle). For each assay, the gel grid was placed on the heated ITO glass and allowed to reach a surface temperature of 35°C, confirmed with thermal camera, before the cage-grid was aligned on top and video recording initiated. Mosquito interactions were additionally recorded by visual inspection to provide redundancy and to guide selection of gel samples and negative controls (gels with confirmed lack of interaction). Assays ended after cessation of activity or after 30 min, whichever occurred first. Gel substrates from interacting mosquitoes and negative controls were collected immediately and frozen.

Two time-course implementations were used:

## Continuous time-course

Infected mosquitoes were introduced into the cage-grid at 9 days post infection (dpi) and repeatedly presented with the bite substrate every other day (10, 12, 14, 16, and 18 dpi) while being maintained within the SpitGrid between sessions. Between assays, cotton pads soaked in 10% glucose (w/v) were placed on the perforated bottoms to maintain mosquitoes until the evening of the next day, after which sugar pads were replaced with water-soaked cotton pads for overnight starvation before the subsequent exposure. Mosquitoes that died during the experiment were collected and dead individuals were replaced with mosquitoes from the same cohort. At the end of the time course, remaining mosquitoes were frozen and collected for parasite quantification by qPCR.

## Discontinuous time-course

The discontinuous design followed the same assay procedure, but instead of maintaining the same individuals, new infected mosquitoes (from the same cohort) were introduced each experiment day. After exposure, mosquitoes were frozen and collected for parasite quantification. Head and thorax was separated from abdomen to approximate salivary gland parasite load.

## Behavioral experiments

Mosquitoes were added to the individual cages and were left to equilibrate for 3 hours prior to start of the experiments. The artificial bite substrate holder was placed on the heating element and once surface temperature was stable at 35°C, confirmed with thermal camera, recording was started and the mosquito cage was placed on top of the bite substrate. At arbitrary moments (e.g. during feeding, probing, or searching behavior), the cage was lifted off and the bite substrate was exchanged with a new bite substrate holder. After exposure, bite substrates were frozen for DNA extraction. Behavioral annotations of video recordings were performed using BORIS^72^.

### DNA extraction from mosquitoes and gel substrates

Parasite loads from salivary glands, thoraces/whole carcasses, and gel substrates were quantified by qPCR. For discontinuous time-course samples, mosquitoes were frozen after the assay. To approximate salivary gland load, thoraces were separated from abdomens and homogenized (bead beating) in PBS. DNA was extracted using the MagNA Pure LC instrument (Roche) with the MagNA Pure LC DNA Isolation Kit – High Performance (Roche, product no. 03310515001) and eluted in 50 µL. For behavioral experiments mosquitoes were knocked down with CO_2_, immobilized, and salivary glands were dissected, homogenized, and extracted as described above. For the continuous time-course variant where whole-body infection intensity was assessed, whole mosquitoes were homogenized and extracted as described above. DNA was extracted from the artificial bite substrates using the peqGOLD Gel Extraction Kit (VWR) according to the manufacturer’s instructions and eluted in 30 µL.

### Quantification

**Standard curves:** To account for differing sample extraction efficiency, separate standard curves were generated for mosquito-derived samples and bite substrate-derived samples. For mosquito standards, infected salivary glands were homogenized in PBS (glass pestle grinder), sporozoites were manually counted using a hemocytometer, serial dilutions prepared, and processed through the MagNA Pure LC workflow. For gel standards, counted sporozoites were introduced into gel substrates and extracted using the gel extraction workflow. Standards were run alongside experimental samples to estimate sporozoite numbers from Ct values.

## Pf qPCR

*P. falciparum* sporozoites were quantified by probe-based qPCR targeting the mitochondrial COX- 1 gene as previously described^13^. Primer/probe sequences were: forward 5’- CATCAGGAATGTTATTGCTAACAC-3’, reverse 5’-GGATCTCCTGCAAATGTTGGGTC-3’, probe 6FAM-ACCGGTTTTAACTGGAGGAGTA-BHQ1. Reactions used TaqMan Fast Advanced Master Mix (Applied Biosystems), 800 nM primers, 400 nM probe, and 40% of the reaction volume as template DNA. Cycling conditions were 95°C for 120 s followed by 40 cycles of 95°C for 15 s and 60°C for 60 s

## Pv qPCR

*P. vivax* sporozoites were quantified using a standard qPCR assay based on Lamien-Meda et al.^73^ with modified primer sequences (forward 5’-GCCTTGCAATAAATTAATATT-3’, reverse 5’- GCCTGGAGTTCTTTATCTTAGTA-3’). Reactions were done with GoTaq mastermix (Promega) used 800 nM primers and 10% of the reaction volume as template DNA. Cycling conditions were 95°C for 120s followed by 40 cycles of 95°C for 15 s and 60°C for 60 s. As for *P. falciparum*, separate standard curves were generated for mosquito-based and gel-based extractions.

## Data analysis and statistics

All analyses and visualizations were performed in Python using standard scientific libraries. Group comparisons used two-sided Mann–Whitney U tests for continuous outcomes and Fisher’s exact tests for proportions. P-values were adjusted for multiple comparisons using Bonferroni correction. As a summary metric on skewed count data while retaining zero values, a pseudo-geometric median was used (log10(x+1) transform). For the *P. vivax* experiments, due to the large amount of zero expellers, two separate summary metrics were used frequency of expelling and geometric median. For time-course modeling of inoculum size, generalized linear models with a log link were fit using a Tweedie distribution^74^. In the time course experiment the effect of days post infection was represented with a piecewise effect (pre- vs post-12 dpi).

