## Supplementary Information for "Quantifying sporozoite inoculum dynamics with the SpitGrid reveals temporal decay and behavioral determinants of *Plasmodium* transmission"

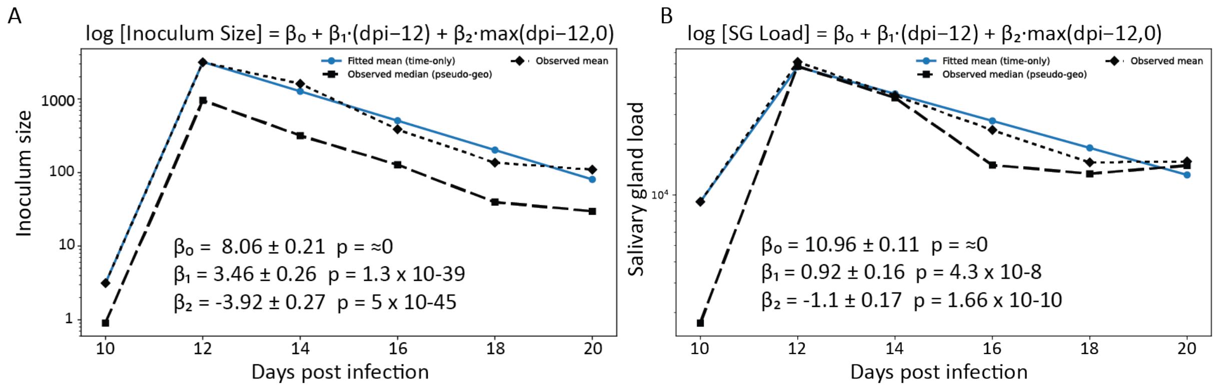


**Supplementary Figure 1.** GLM captures time-dependent expelling trajectory and salivary gland infection. **A** fitted mean inoculum size over time from a broken-stick (knot at 12 dpi) model on log-transformed gel sporozoite counts, shown alongside the observed mean and observed pseudo-geometric median. **B** Same analysis for salivary gland loads. Parameter estimates and Wald-test *p*-values are displayed in-panel.

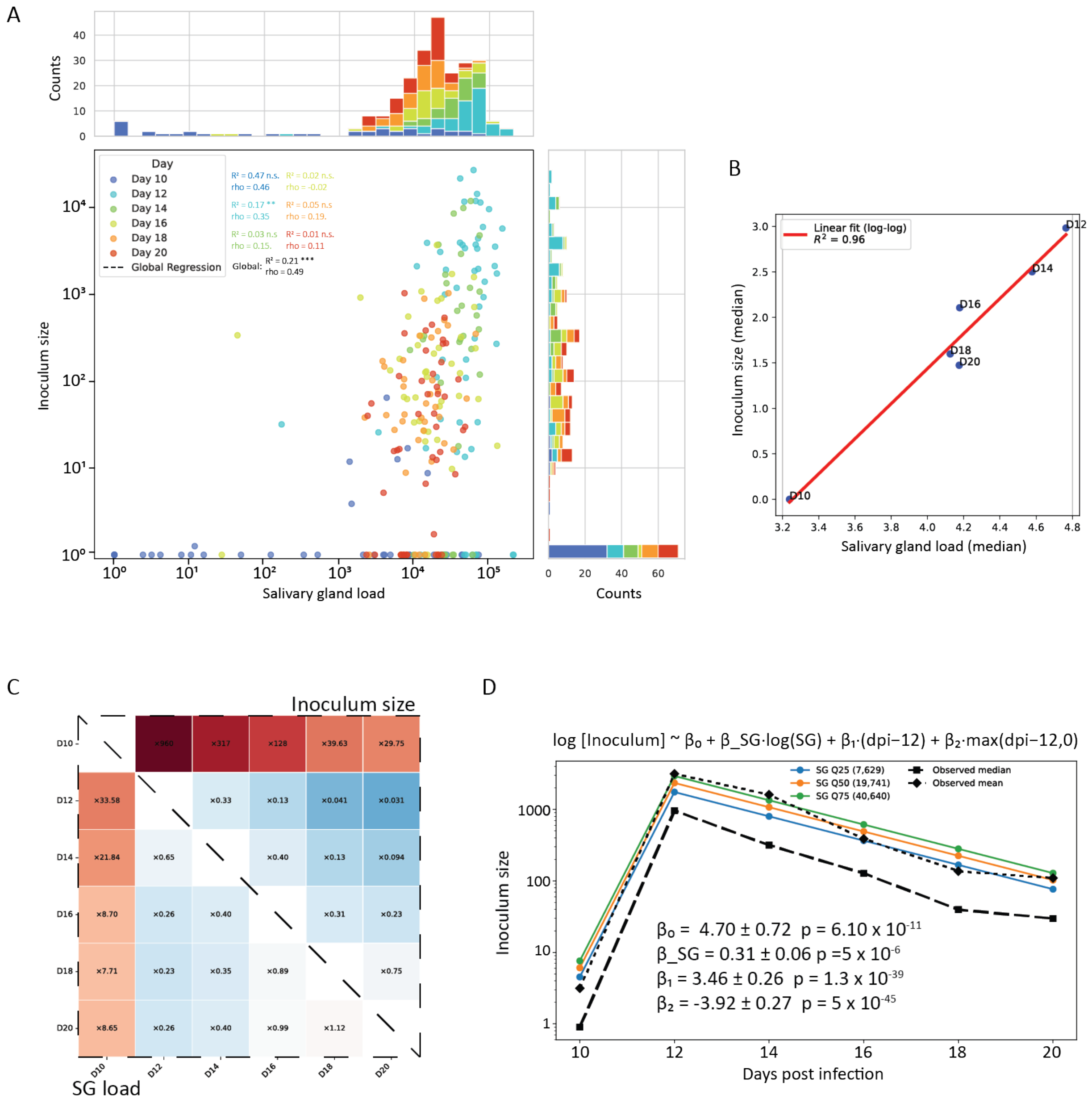


**Supplementary Figure 2.** Relationship between salivary gland load and inoculum size across days post infection. **A** Scatter plot of inoculum size versus salivary gland load for individual mosquitoes, colored by dpi, with marginal histograms and per-day correlation statistics and a global regression across all days are displayed in panel. **B** Association between day-level medians of salivary gland load and inoculum size in log–log space with linear fit and $R^2$. **C** Pairwise fold-change matrix across dpi for salivary gland load (lower triangle) and inoculum size (upper triangle), visualizing day-to-day shifts in median values. **D** Time model for inoculum size including salivary gland load as covariate, showing fitted mean trajectories at representative SG-load quantiles (with observed mean/median overlaid); model coefficients and p-values are shown in-panel.

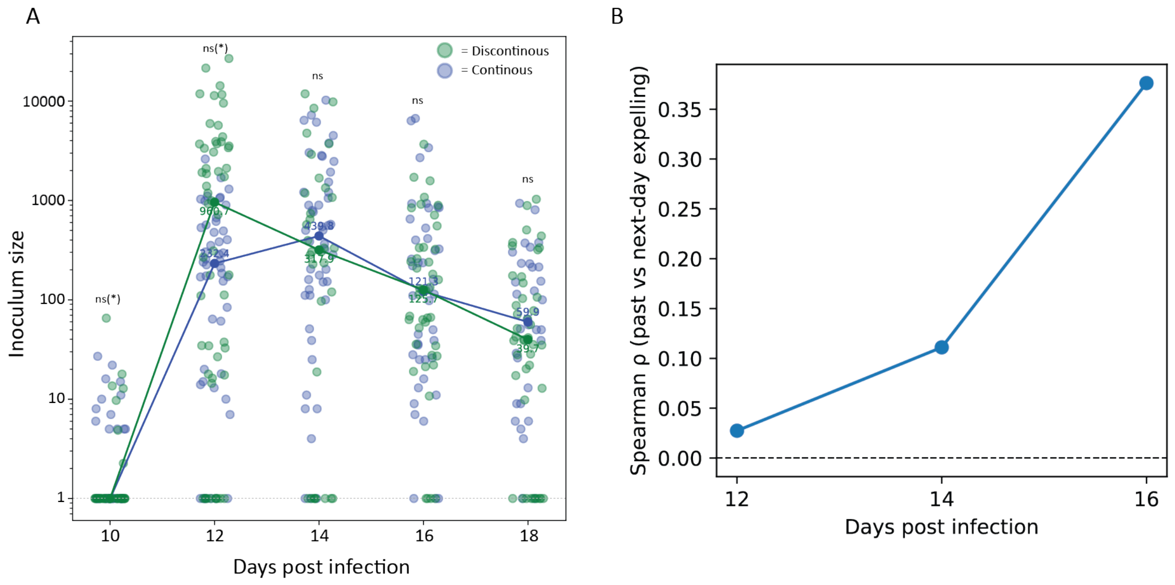


**Supplementary Figure 3.** Continuous and discontinuous timecourses show similar inoculum dynamics and increasing within-mosquito consistency over time. **A** Inoculum size distributions across dpi for discontinuous and continuous experiments with per-day pseudo-geometric medians overlaid; in panel p-values above days indicate between-experiment comparisons (two-sided MWU, uncorrected). **B** Spearman correlation (ρ) between sporozoites expelled on a given day and the subsequent exposure day within the continuous timecourse experiment.

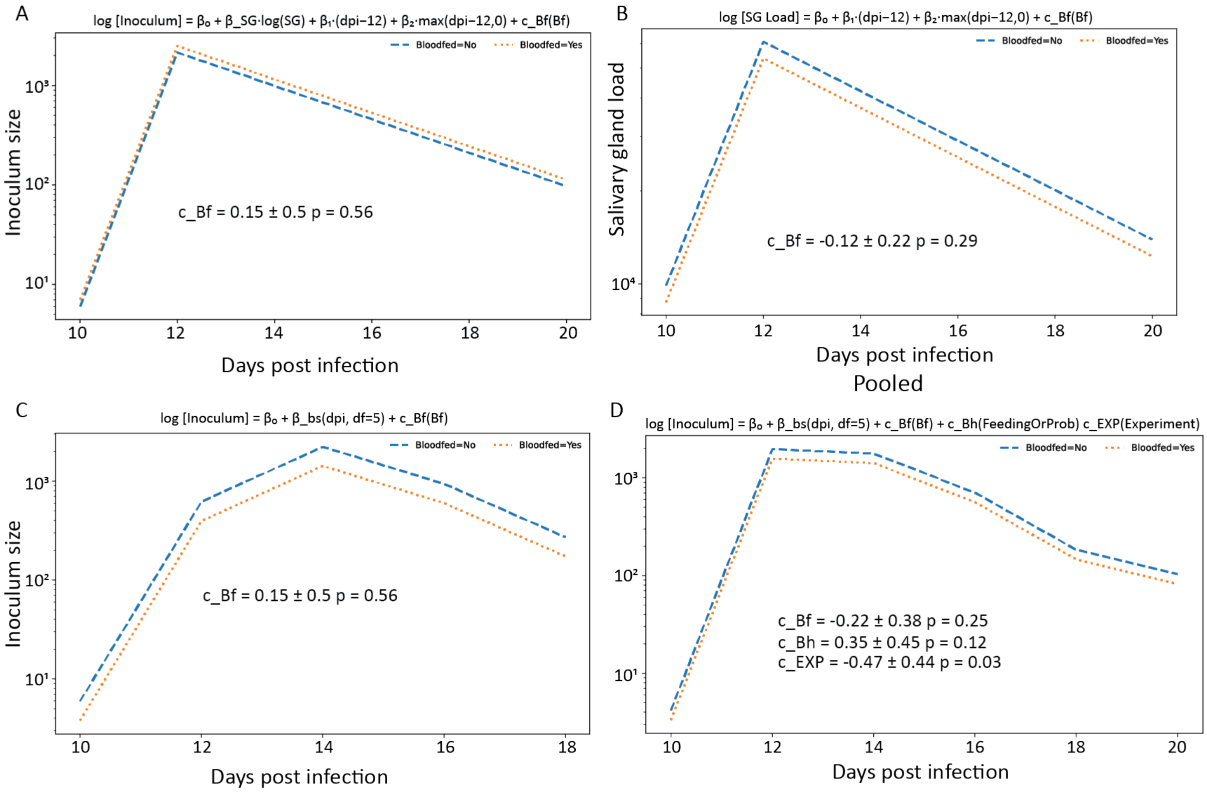


**Supplementary Figure 4.** A second non-infectious bloodmeal does not measurably shift inoculum size or salivary gland loads in regression models. **A** Relative frequency of salivary gland load bins (top) and inoculum-size bins (bottom) across dpi for mosquitoes receiving only the infectious bloodmeal versus an additional non-infectious bloodmeal. **B** Discontinuous timecourse GLM for inoculum size including SG load and broken-stick time terms, with an added bloodfed term; fitted mean trajectories for bloodfed versus non-bloodfed groups are shown and the estimated bloodfed effect (c_Bf) is annotated in panel. **C** Corresponding GLM for SG load with a bloodfed term. **D** Continuous timecourse GLM for inoculum size with bloodfed term only (time modeled with a spline/basis as indicated), showing fitted trajectories and c_Bf estimate in panel. **E** Pooled model additionally including interaction type (feeding vs probing-only) and experiment indicator; corresponding coefficients and Wald p-values are shown in panel.

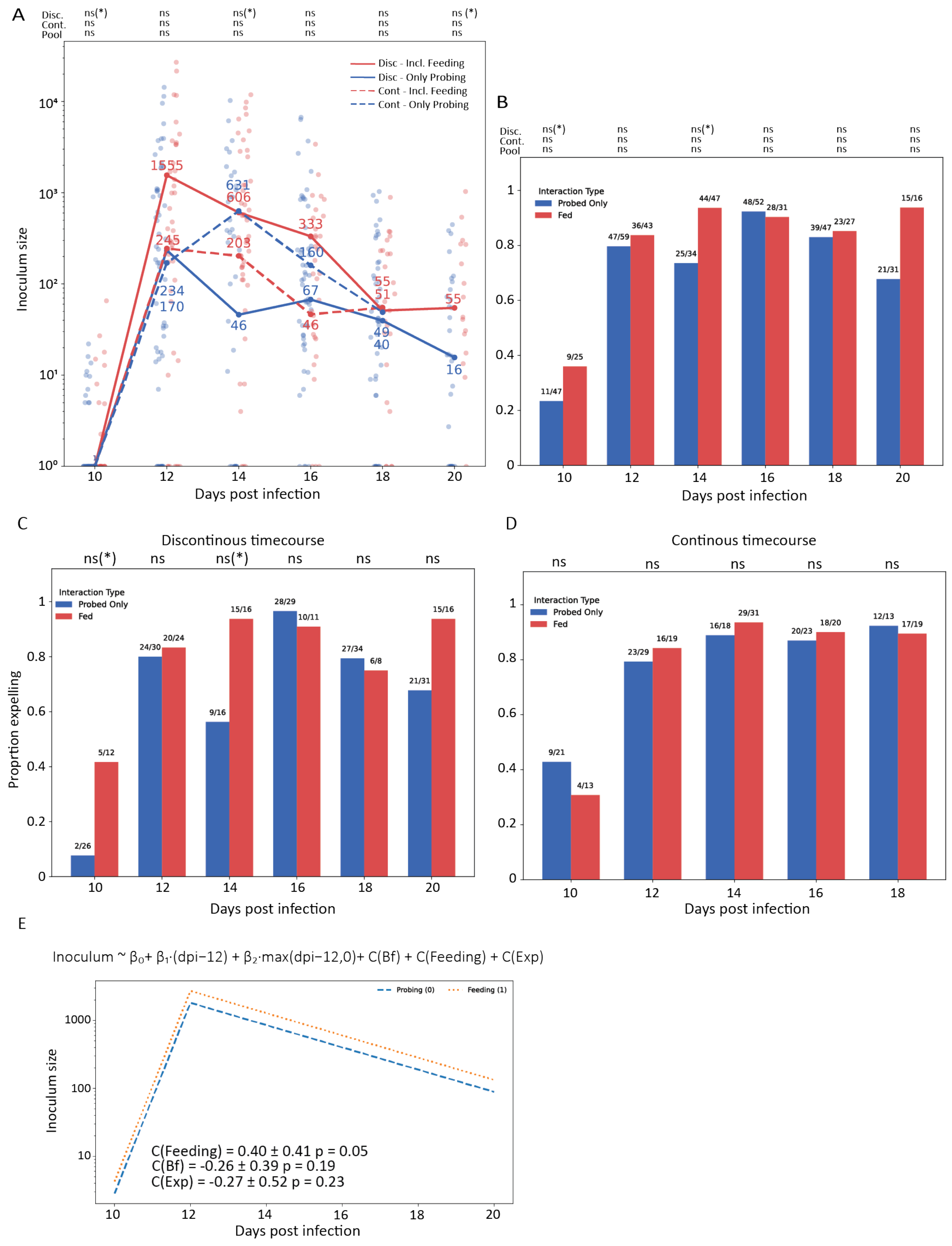


**Supplementary Figure 5.** Probing-only interactions yield similar inoculum sizes and expelling prevalence as interactions that include feeding. **A** Inoculum size over time stratified by interaction type (feeding vs probing-only) and experiment implementation (continuous vs discontinuous), with pseudo-geometric medians annotated; significance summaries above indicate per-day comparisons within each experiment and pooled. Brackets denote significance prior to correction **B** Proportion of mosquitoes expelling sporozoites (bite substrate positive) by interaction type across dpi (pooled), with absolute counts shown above bars and per-day significance annotations. Brackets denote significance prior to correction. **C** Same as (B) for the discontinuous experiment only. **D** Same as (B) for the continuous experiment only. **E** Pooled GLM of inoculum size including broken-stick time terms and categorical predictors for bloodfed status, interaction type, and experiment; fitted mean trajectories for probing-only vs feeding are shown with coefficient estimates and Wald p-values in panel.

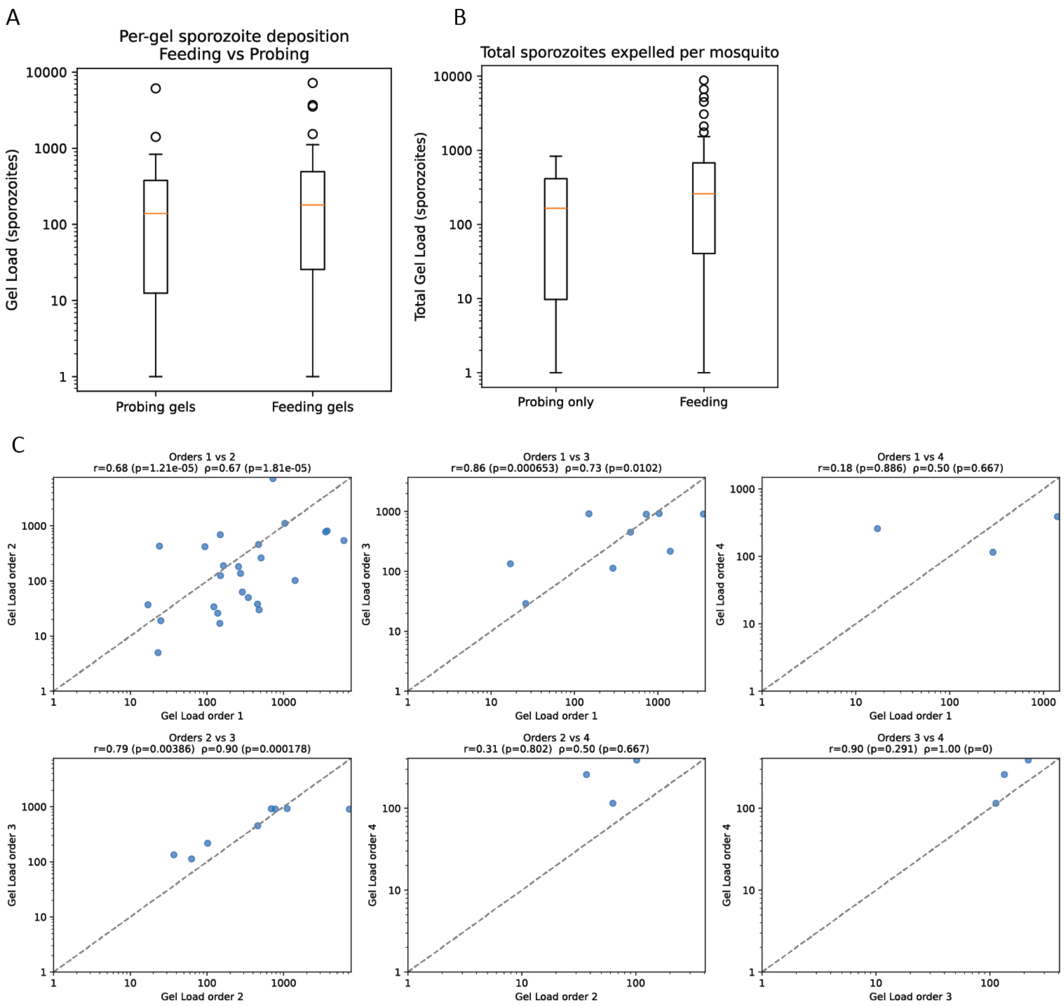


**Supplementary Figure 6.** High-resolution behavior analysis shows that feeding does not increase per-gel or per-mosquito sporozoite deposition, and successive gels are correlated. **A** Per-gel sporozoite deposition comparing gels from probing-only interactions versus interactions that included feeding (boxplots on log scale). **B** Total sporozoites expelled per mosquito (summed across gels) comparing mosquitoes that only probed versus those that fed at least once (boxplots on log scale). **C** Pairwise comparisons of inoculum size between successive gel orders within mosquitoes (orders 1–4), showing Pearson r and Spearman ρ with p-values; dashed diagonal indicates y=x.

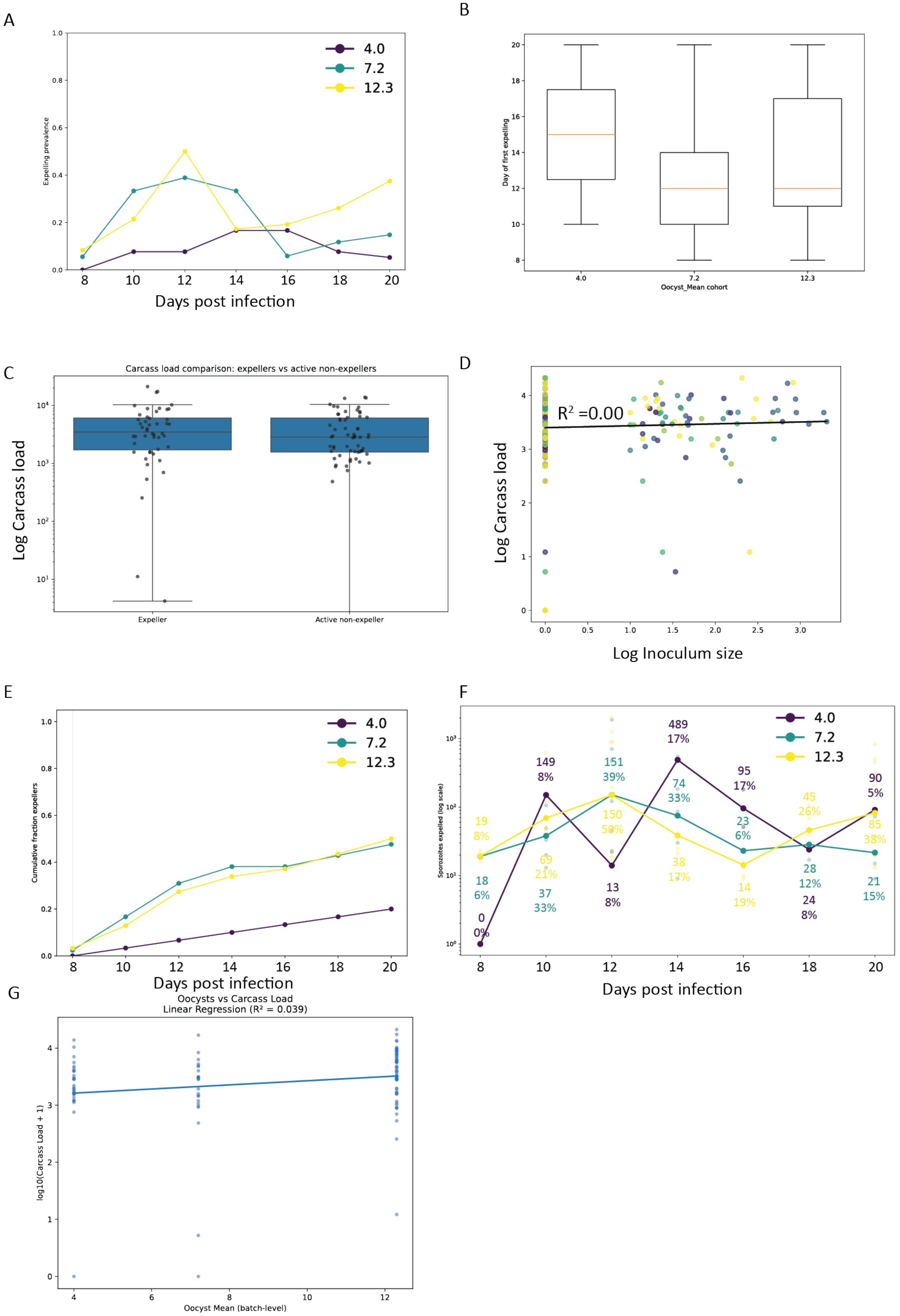
A
Expelling prevalence
Days post infection
4.0
7.2
12.3
B
Day of first expelling
Oocyst_Mean cohort
C
Carcass load comparison: expellers vs active non-expellers
Log Carcass load
Expeller
Active non-expeller
D
$R^2$ =0.00
Log Carcass load
Log Inoculum size
E
Cumulative fraction expellers
Days post infection
F
Sporozoites expelled (log scale)
Days post infection
G
Oocysts vs Carcass Load
Linear Regression ($R^2$ = 0.039)
log10(Carcass Load + 1)
Oocyst Mean (batch-level)

**Supplementary Figure 7.** *P. vivax* cohort-level oocyst intensity relates to expelling prevalence, while carcass load does not explain inoculum variation. **A** Per-day expelling prevalence (fraction of interacting mosquitoes with detectable sporozoites) stratified by cohort mean oocyst intensity. **B** Distribution of time (dpi) to first detectable expelling by cohort (boxplots). **C** Carcass parasite load comparing mosquitoes that expelled at least once versus “active non-expellers” (mosquitoes that interacted but never expelled detectable sporozoites). **D** Association between log inoculum size and log carcass load with linear fit and $R^2$ annotation. **E** Cumulative fraction of mosquitoes that have expelled at least once over time, stratified by oocyst cohort. **F** Inoculum size over time (log scale) stratified by cohort with day-level geometric medians annotated and corresponding daily expelling rates annotated above. **G** Relationship between cohort mean oocyst intensity (batch-level) and individual carcass load with linear regression and $R^2$ annotation.

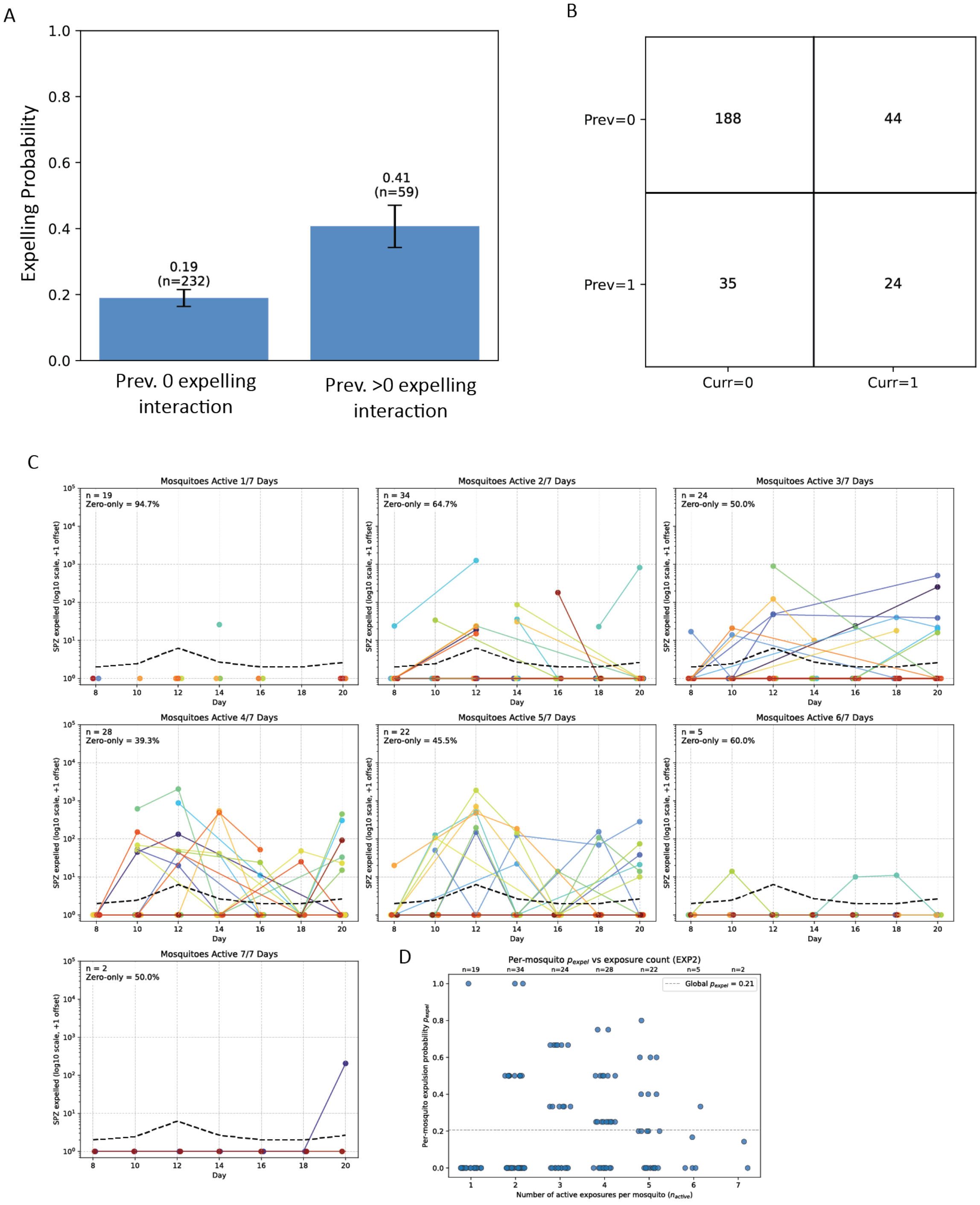


**Supplementary Figure 8.** *P. vivax* expelling events are not independent and mosquitoes differ in expelling propensity. **A** Conditional probability of expelling on the current exposure given whether the previous exposure had zero versus >0 expelled sporozoites. Bars show estimated probabilities with error bars and sample sizes. **B** Matrix of previous (Prev) versus current (Curr) expelling outcome counts. **C** Individual mosquito inoculum trajectories grouped by the number of active days (days with probing/feeding), highlighting frequent zero-expellers despite interaction. Dashed line shows the mean across days. **D** Per-mosquito expelling probability (fraction of active exposures with detectable sporozoites) plotted against the number of active exposures, with the global average indicated.

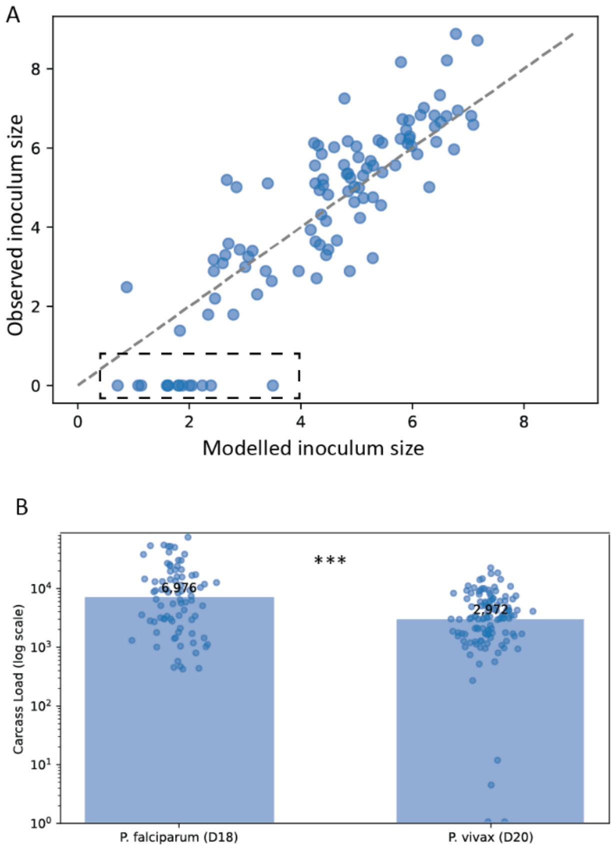


**Supplementary Figure 9.** Additional analyses. **A** Model evaluation plot comparing modelled versus observed inoculum size for individual observations; dashed line indicates y=x and the boxed region highlights prediction error in low/zero-inoculum observations. **B** Comparison of terminal carcass parasite loads between P. falciparum (D18) and P. vivax (D20) with individual datapoints overlaid and group medians annotated; significance is indicated above the comparison.
